# Multi-Omics Single-Cell Analysis Reveals Key Regulators of HIV-1 Persistence and Aberrant Host Immune Responses in Early Infection

**DOI:** 10.1101/2024.11.04.621999

**Authors:** Dayeon Lee, Sin Young Choi, So-I Shin, Hyunsu An, Byeong-Sun Choi, Jihwan Park

**Affiliations:** School of Life Sciences, Gwangju Institute of Science and Technology (GIST); Gwangju, Republic of Korea; Director for Laboratory Diagnosis and Analysis Chungcheong Regional Center for Disease Control and Prevention Korea Disease Control and Prevention Agency (KDCA); 35208, Republic of Korea

**Keywords:** HIV-1, CD4+ T cell, Acute infection, single-cell RNA sequencing, single-cell ATAC sequencing

## Abstract

The clearance of human immunodeficiency virus-1 (HIV-1) remains a significant public health challenge due to impaired cellular immune responses and HIV-1 maintenance during acute infection. However, the genetic and epigenetic changes influencing the immune response on host infected cells remain unclear. Here, this study analyzes HIV-1 infected CD4+ T cells from peripheral blood mononuclear cells from people living with HIV-1 (PLWH) during early infection (<6 months) using single-cell RNA and ATAC sequencing. It is observed that HIV-1 hinders the antiviral response, particularly by interfering with the interferon signaling pathway. Multimodal analysis identifies KLF2 as a key transcription factor in infected CD4+ T cells. Moreover, cells harboring HIV-1 provirus are predominantly identified as Th17 cells, which exhibit elevated KLF2 activity. This suggests an increased susceptibility to HIV-1 infection and a constrained immune response due to the quiescent characteristics of these cells. The finding provides insights into the immune mechanisms and key regulators of HIV-1 maintenance in CD4+ T cells during the early stages of infection.

## INTRODUCTION

Each year, approximately two million people are infected with human immunodeficiency virus-1 (HIV-1), and about 1.1 million individuals die from HIV-related illnesses. HIV-1 primarily targets CD4+ T lymphocytes, leading to a gradual depletion of this critical T-cell population and the eventual onset of acquired immunodeficiency syndrome (AIDS) [1, 2]. Although antiretroviral therapy (ART) helps inhibit HIV-1 replication [3], the persistent presence of the virus causes chronic inflammation, contributing to HIV-related morbidity and mortality [4, 5]. Additionally, latent proviruses can be reactivated and causing viral rebound if ART is discontinued. As a result, achieving complete clearance of HIV-1 remains a major goal and public health challenge [6].

During HIV-1 infection, cellular immune responses are disrupted, leading to immune cell dysfunction and alterations in the immune cell landscape. Acute HIV infection (AHI) occurs from 3 weeks to 6 months after transmission, following an initial period when the viral load is undetectable [7]. Once proviruses integrate into the host genome, they impact the recruitment of regulatory factors, thereby affecting gene expression [8–10]. These modifications during AHI play a role in the host cell’s initial immune response and contribute to HIV-1 latency [6, 11, 12]. Studying transcriptomic changes during AHI is crucial to identify primary immune responses and regulatory factors that influence HIV-1’s effect on host immune cell function.

Single-cell RNA sequencing (scRNA-seq) and single-cell Assay for Transposase Accessible Chromatin with high-throughput sequencing (scATAC-seq) are powerful techniques that provide insights into cellular diversity, allowing for high-resolution analysis of gene expression and regulatory landscapes in individual cells [13–18]. Recent advancements in sequencing technologies have driven numerous studies to uncover biological factors contributing to HIV-1 pathogenesis [19–24]. However, challenges remain in studying genetic and epigenetic regulation in HIV-1-infected cells: low HIV-1+ CD4+ T cell frequency in PBMCs limits reliable analysis [21, 25], ex vivo activation in previous studies does not capture in vivo heterogeneity [20, 26, 27]. Additionally, although recent studies have attempted direct ex vivo omics analyses [28–30], integrating these datasets remains difficult due to the varying cell types used in each dataset.

To address these limitations, we conducted single-cell multi-omics analyses on PBMCs from people living with HIV (PLWH) in the early stages of infection. Using a targeted sequencing strategy, we identified and analyzed HIV-1-infected cells at the single-cell level. Based on these findings, we established a gene regulatory network specific to HIV-1-infected cells. This study highlights multiple immune pathways and regulatory factors mediating the effects of acute HIV-1 infection on immune cell function, offering potential therapeutic targets for complete HIV-1 eradication.

## RESULTS

### The single cell transcriptional landscape of HIV-1 infected cells in early stage of infection

To uncover immune mechanisms and transcriptional regulation in HIV-1-infected cells, we performed single-cell multiomics sequencing on PBMCs from nine individuals with early-stage HIV infection (< 6 months) (Supplementary Table 1). Using the 10x Genomics Single-Cell platform, we obtained five scRNA-seq and four snRNA-seq data from single cell multiome datasets. We integrated scRNA-seq and snRNA-seq data, identifying 16 distinct cell populations based on marker genes (Fig. 1A, Supplementary Fig. 1A). We also observed some variability in the proportions of these cell populations among donors, suggesting inter-individual differences in immune cell composition (Supplementary Fig. 1B). Focusing on CD4+ T cells as main targets of HIV-1, we defined four subtypes: CD4 naïve T cells, CD4+ effector memory T cells (CD4 EM cells), T helper (Th)2 cells, and Th17 cells (Fig. 1B, Supplementary Fig. 1C, 1D). *LDHB* and *SELL* were highly expressed in CD4 naïve T cells, whereas CD4 EM cells were characterized by high expression levels of *HLA-DRA* and *GZMK*. Among the activated CD4+ T cells, Th2 cells were identified based on the expression of *STAT6* and *GATA3* and Th17 cells were annotated based on the expression levels of *RORA* and *STAT3*.

**Fig. 1.**
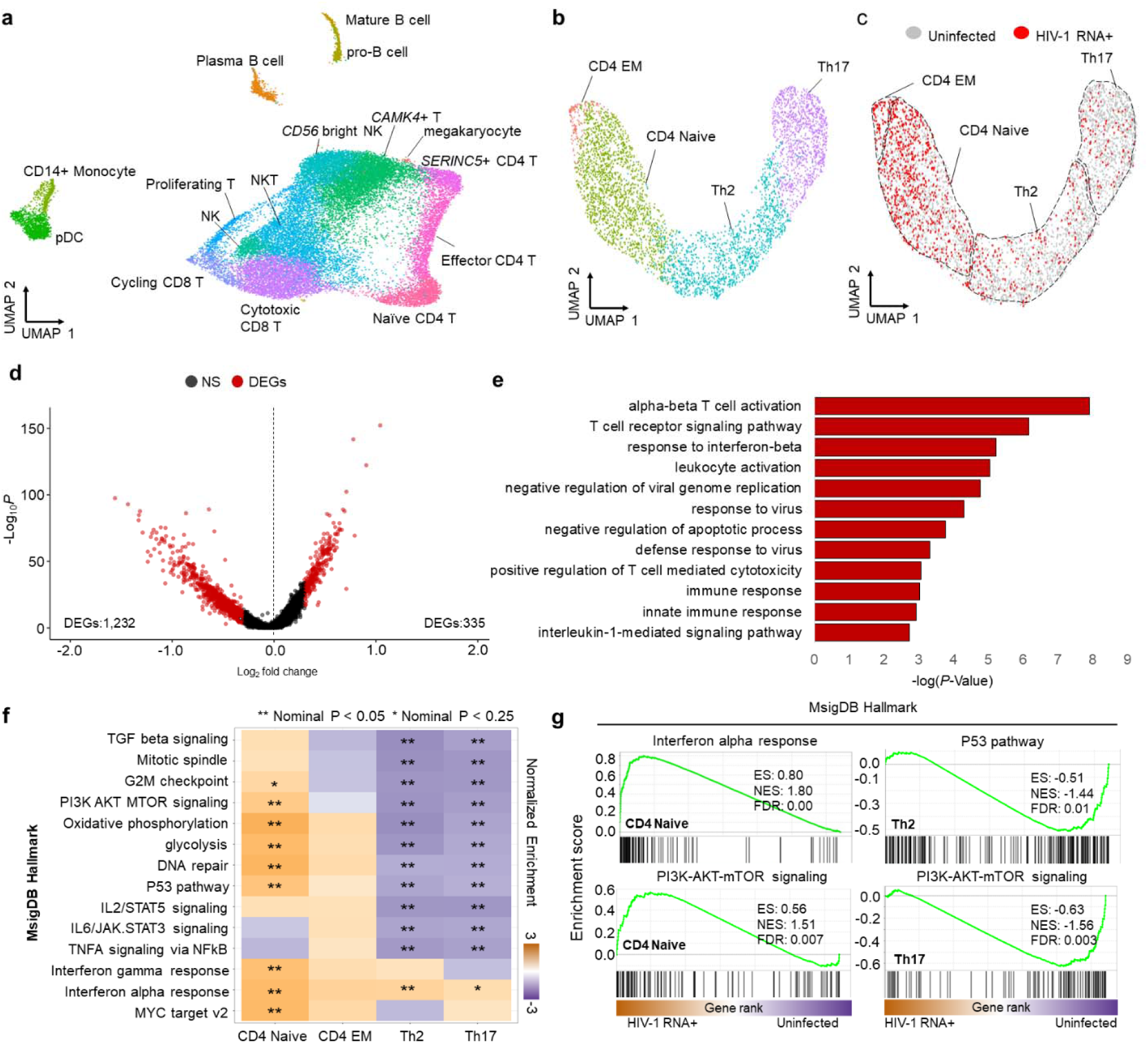
Single Cell Transcriptomic Analysis of HIV-1 RNA+ cells from Early Infected Patients. (a) UMAP plot displays the distribution of PBMCs in early infected patients (n=9), clustered based on transcriptome signatures post-removing batch effects. (b) The UMAP plot displays the distribution of 4,435 CD4 T cells from early infected patients. (c) UMAP of CD4 T cells identifies 3,450 uninfected cells and 985 HIV-1 RNA+ cells. (d) Volcano plot reveals differentially expressed genes (DEGs) in HIV-1 RNA+ CD4 T cells compared to uninfected cells; red dots signify significant DEGs (adj *P*-value < 0.01). (e) Bar plot reveals enriched biological processes and immune pathways in HIV-1 RNA+ CD4 T cells (*P*-value < 0.01) using upregulated DEGs; significance presented as −log(*P*-value). (f) Heatmap exhibits signaling pathways enriched in CD4 T cell types through GSEA; positive scores (dark orange) denote enrichment in HIV-1 RNA+ cells (Nominal *P*-val < 0.05 and < 0.25) are marked with stars. (g) GSEA plots show significant immune pathways in CD4 naïve, Th2 and Th17 cells, presenting enrichment score (ES), normalized enrichment score (NES), and false discovery rate (FDR).

To identify HIV-1 RNA+ cells, we aligned scRNA-seq reads to the HIV-1 genome. Additionally, we conducted targeted long-read sequencing using the remaining scRNA-seq libraries to improve the sequencing depth of HIV-1 transcripts. We identified 985 HIV-1 RNA+ cells and 3,450 uninfected cells. CD4 naïve and EM cells showed a higher proportion of HIV-1 RNA+ cells than Th2 and Th17 (Fig. 1C, Supplementary Fig. 1E).

Gene expression profiling revealed 334 upregulated and 1,002 downregulated genes in HIV-1 RNA+ CD4+ T cells (Fig. 1D), including genes related to T cell activation (*PRDX2*, *PRDX1*, *HLA-B*, *HLA-C*, *HLA-A*, *HLA-F*, and *HLA-E),* defense responses to viruses (*IFITM3*, *IFITM1*, *IFITM2*, and *MX1*), and interferon-beta responses (*IFITM3*, *BST2*, *IFITM1*, *IFITM2*, *XAF1*, and *SHFL*) (Fig. 1E). Notably, IFITM1 was the most upregulated gene in HIV-1 RNA+ CD4+ T cells. As an interferon-stimulated gene, IFITM1 is known to inhibit HIV-1 replication by interfering with viral entry [31]. This finding indicates that the primary cellular pathways active during acute infection are also engaged in the antiviral response to suppress HIV-1.

Given that CD4+ T cell differentiation influences HIV-1 survival [32], we used gene set enrichment analysis (GSEA) to assess subtype-specific responses (Fig. 1F). The GSEA results revealed that interferon-gamma and interferon-alpha responses were upregulated in all CD4+ T-cell subtypes upon HIV-1 infection. Additionally, hallmarks related to T-cell activation were downregulated in both Th2 and Th17 cells. Additionally, while interferon-alpha responses were enriched, downstream pathways (e.g., PI3-AKT-mTOR) were downregulated in Th17 cells (Fig. 1F, G). Downregulated genes in Th17 cells were linked to PI3K-AKT signaling, while no such enrichment was observed among upregulated genes (Supplementary Fig. 2A, 2B). These results suggest that activated T cells experience immune dysfunction in early HIV-1 infection, contrasting with the effects observed in naïve CD4+ T cells.

### Characterization of epigenetic changes in HIV-1 infected CD4+ T cells in early infection

Single-cell multiome technology enables a detailed understanding of cell states by simultaneously profiling transcriptome and epigenetic signatures within the same cell, facilitating the construction of an accurate gene regulatory network by identifying gene expression drivers. A combined UMAP was generated based on chromatin accessibility and gene expression, with cell types annotated using marker genes (Fig. 2A, Supplementary Fig. 3A). Additionally, coverage plots revealed cell type-specific chromatin accessibility for nearby marker genes (Fig. 2B). Chromatin accessibility in HIV-1 RNA+ naïve and memory CD4+ T cells identified specific patterns, with closed DARs in naïve cells linked to TNF-alpha signaling via NF-κB and open DARs related to HIV-1 Nef protein and signal transduction. In memory CD4+ T cells, closed DARs were related to interferon signaling and negative regulation of viral transcription, whereas open DARs were related to Tat-mediated HIV elongation. When examining genes adjacent to open chromatin regions (Fig. 2D), we observed high chromatin accessibility for *PSMB7* and *CDK9* in naïve and memory CD4+ T cells, respectively, both of which interact with the HIV-1 Tat protein [33, 34]. Furthermore, the chromatin accessibility of *NR4A2* and *TRIM62*, which negatively regulate HIV-1 [35–37], was reduced in HIV-1 RNA+ cells (Fig. 2E).

**Fig. 2.**
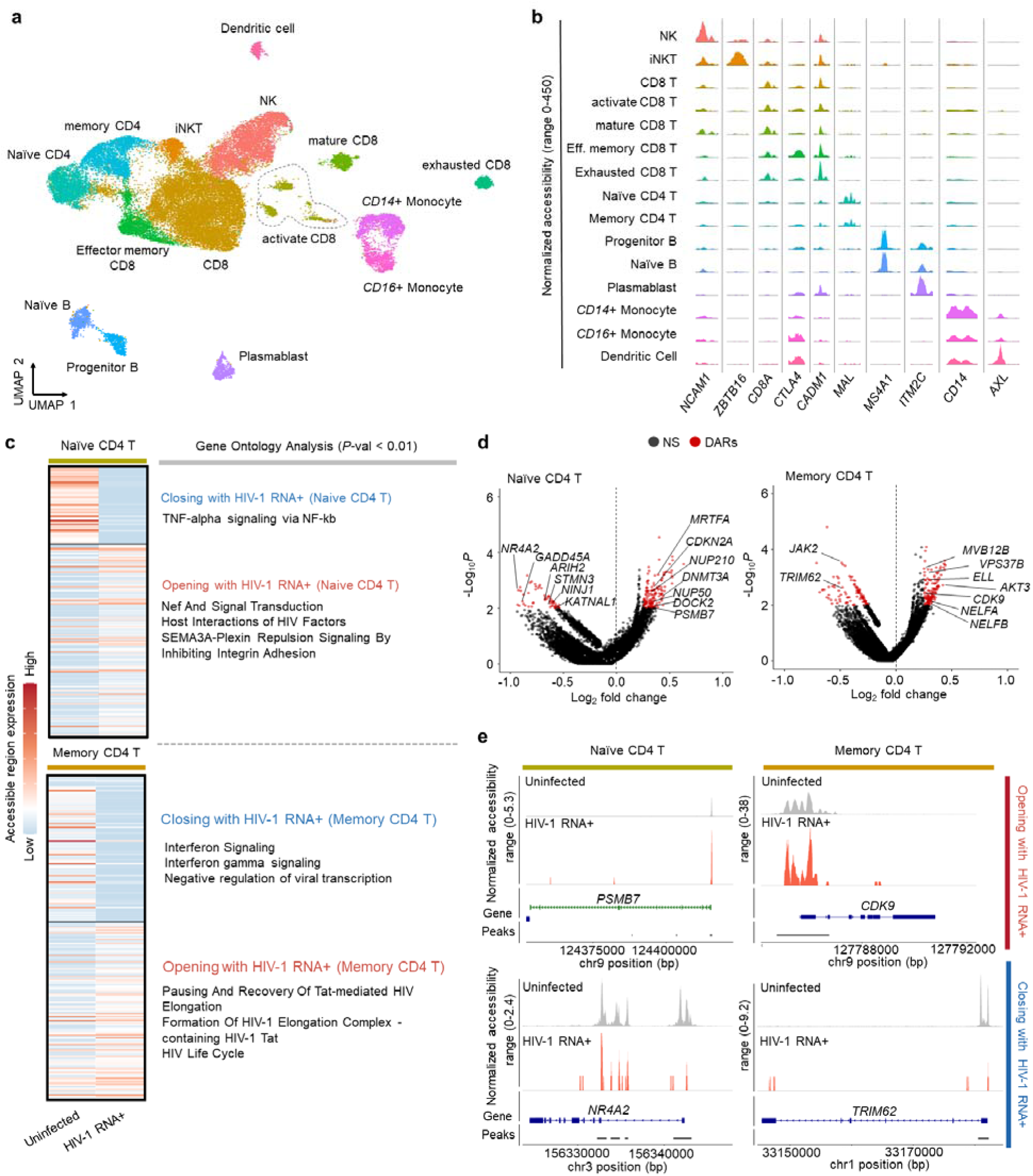
Exploring The Epigenetic Characteristics of HIV-1 RNA+ Cells Through Single-Cell Multiome Data. (a) UMAP plot illustrates PBMC distribution in early infected patients (n=8) post-clustering based on transcriptome and epigenome signatures, with batch effects removed. (b) Visualization of differential DNA accessibility across 15 cell clusters. Tracks display normalized chromatin accessibility at promoter regions of cluster-specific marker genes. (c) Heatmap displays differentially chromatin accessible regions (DARs) in HIV-1 RNA+ and uninfected cells of Naïve and Memory CD4 T cell types. Right heatmap presents enriched gene ontology for DAR-associated genes. (d) Volcano plot shows DARs between HIV-1 RNA+ CD4 T cells and uninfected cells; red dots signify significant DARs, with associated genes displayed. (e) Tracks display normalized chromatin accessibility at the promoter regions of key genes (*PSMB7*, *CDK9*, *NR4A2*, *TRIM62*) in HIV-1 RNA+ and uninfected cells.

### Identification of key transcription factors (TFs) governing HIV-1 infection features

Following HIV-1 infection, host transcription factors (TFs) regulate immune responses and viral persistence [38]. To identify key TFs and construct a gene regulatory network associated with HIV-1 infection, we integrated gene expression and chromatin accessibility data from HIV-1 RNA+ CD4+ T cells.

We found 15 TFs with increased and 24 with decreased regulon activity in HIV-1 RNA+ cells compared to uninfected cells across all CD4+ T-cell subtypes (Fig. 3A). We also confirmed the significantly different regulon activity of these TFs by cell type. CD4 naïve and EM cells shared similar TF activity profiles, while Th17 and Th2 cells exhibited more comparable patterns (Fig. 3B). KLF2, typically high in naïve cells and suppressed in activated cells [39–41], showed elevated regulon activity in naïve and EM cells, with lower activity in Th17 and Th2 cells (Fig. 3B). However, in HIV-1-infected cells, KLF2 activity was upregulated across all subtypes (Fig. 3A). Conversely, FOXO1 showed higher regulon activity in Th17 and Th2 cells but was downregulated in HIV-1 RNA+ Th17 and Th2 cells. In HIV RNA+ cells, the TFs KLF2 and JUND showed strong regulatory activity and elevated gene expression (Fig. 3C). Additionally, we identified the upregulated target genes of six TFs (KLF2, JUND, STAT1, JUNB, XBP1, MYC) in HIV-1 RNA+ cells (Fig. 3D), which were significantly associated with the defense response to viruses, negative regulation of viral genome replication, and innate immune responses (Supplementary Fig. 3B). As the GSEA result showed different enriched signaling pathways in HIV-1 RNA+ cells by various cell types (Fig. 1G), several key transcription factor-encoding genes (*KLF2*, *FOXO1*, *TBL1XR1*) which regulate PI3K-ATK signaling and p53 signaling, were also differentially expressed in each cell types. (Fig. 3E)

**Fig. 3.**
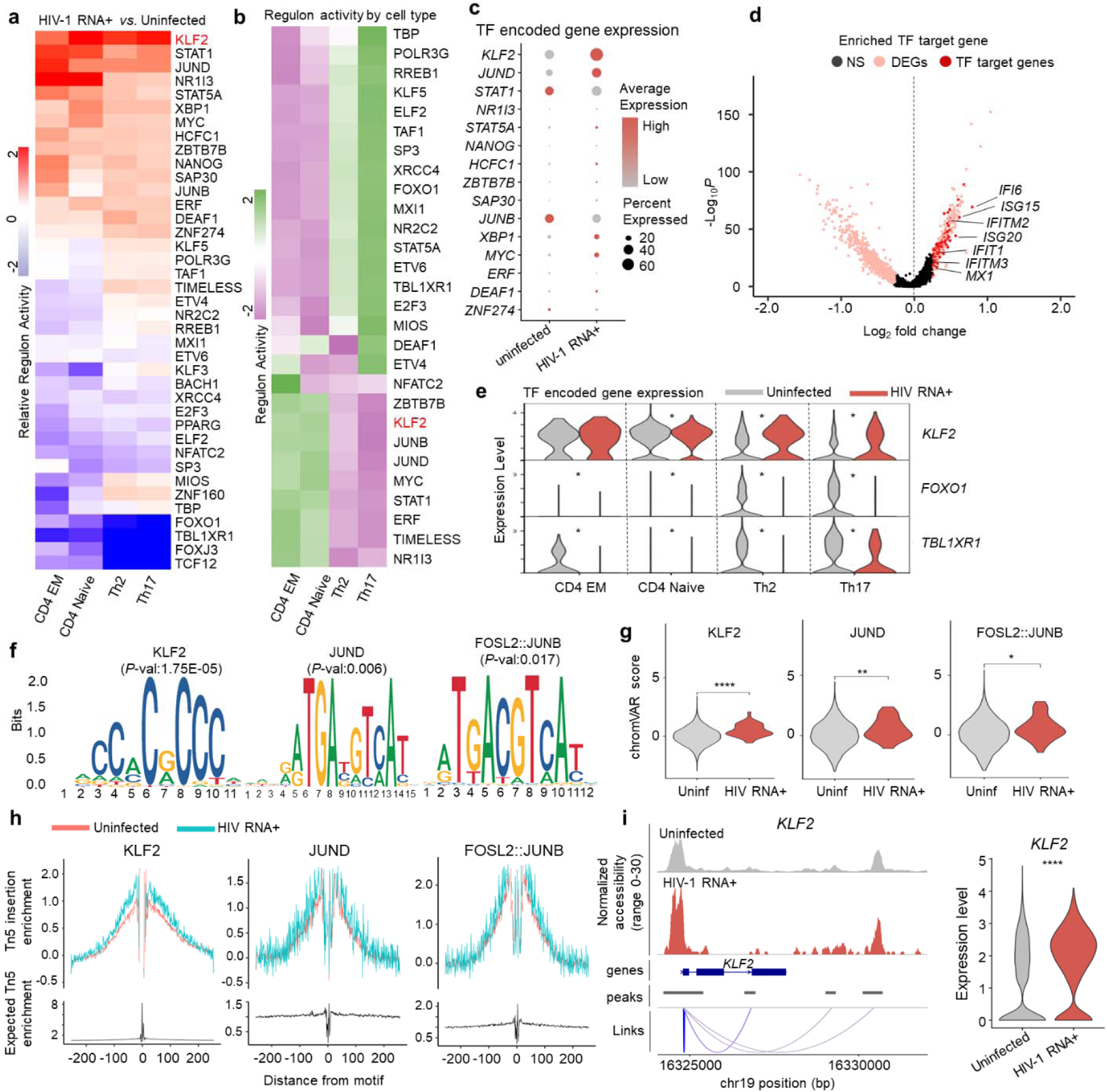
Identifying Key Regulators of HIV-1 RNA+ CD4 T Cells Through Integrated Analysis. (a) Heatmap depicts relative regulon activity in each CD4 T cell subtype between HIV-1 RNA+ and uninfected cells. Positive values indicate increased activity in HIV-1 RNA+ cells, while negative values represent a decrease. (b) Heatmap displays regulon activity levels in each CD4 T cell subtype, with color indicating scaled regulon activity. (c) Dot plot displays gene expression of upregulated TFs in HIV-1 RNA+ cells, with dot size representing the percentage of cells expressing the gene. (d) Volcano plot reveals target genes of upregulated TFs in HIV-1 RNA+ cells; pink dots represent DEGs, and red dots indicate target genes of enriched TFs. (e) Violin plot illustrates expression of TF-encoding genes in uninfected and HIV-1 RNA+ CD4 T cells across subtypes (\**P* < 0.01). (f) DNA sequences for overrepresented TF binding motifs in HIV-1 RNA+ T cells compared to uninfected cells. (g) Violin plot displays chromVAR motif activity score for enriched motifs. (h) TF footprinting profiles show the increased chromatin accessibility near three representative motifs in HIV-1 RNA+ cells. (i) Left panel: Tracks display normalized chromatin accessibility at KLF2 gene locus for HIV-1 RNA+ and uninfected cells, with peak-to-gene links bottom of the coverage plot. Right panel: Expression level of KLF2 in each group.

The chromatin accessibility data further supported the increased activity of these TFs in HIV-1 RNA+ cells. We calculated TF activity based on chromatin accessibility and identified enriched TF-binding motifs in HIV-1 RNA+ CD4+ T cells using DARs. KLF2-, JUND-, and FOSL2::JUNB-binding motifs were highly enriched in the DARs opened in HIV-1 RNA+ cells (Fig. 3F), along with enhanced TF activity (Fig. 3G). Furthermore, TF footprinting analysis revealed higher chromatin accessibility adjacent to the KLF2-binding motif sites (Fig. 3H). The promoter region of *KLF2* was more accessible, with higher mRNA expression in HIV-1 RNA+ cells than in uninfected cells (Fig. 3I). Furthermore, the putative cis-regulatory element of KLF2 showed more chromatin-accessible peaks in HIV-1 RNA+ CD4+ T cells than in uninfected cells (Fig. 3I). Overall, this multidimensional approach enabled integrating gene expression and chromatin accessibility in the early stage of infection in PLWH, highlighting potential key TFs that regulate HIV-1 transcription in host cells.

### KLF2 is an essential regulator in the function of HIV-1-infected CD4+ T cells

Using a multidimensional approach, we identified KLF2 as a key TF within the gene regulatory network of HIV-1 RNA+ CD4+ T cells. KLF2 is reported to be involved in the quiescence, survival, and differentiation of T cells [42]. However, its function in viral infection remains poorly understood. To elucidate the role of KLF2, we curated 88 KLF2 target genes from published chromatin immunoprecipitation (ChIP)-chip, ChIP-seq, and other TF-binding site profiling data, as well as SCENIC regulon datasets. Out of 88 KLF2 target genes, 44 were upregulated in HIV-1 RNA+ cells (Fig. 4A, Supplementary Table 2). To validate these findings, we analyzed a publicly available single-cell transcriptomic dataset of HIV-infected PBMCs [43]. Consistently, expression of *KLF2* and a significant proportion of KLF2 target genes, including *LTB*, *PIM1* and *NRP2*, were also upregulated in HIV-infected CD4+ T cells (Supplementary Fig. 4A). These target genes were mainly associated with negative regulation of apoptotic process and negative regulation of programmed cell death (Fig. 4B). Among these significantly upregulated KLF2 target genes, those reported to be involved in the regulation of HIV-1 replication (*PIM1* and *IL32*) [44, 45], cell proliferation (*TXNIP*) [46], and the JAK/STAT pathway (*LTB*) also exhibited higher levels of chromatin accessibility in HIV-1 RNA+ CD4+ T cells than in uninfected cells (Fig. 4C).

**Fig. 4.**
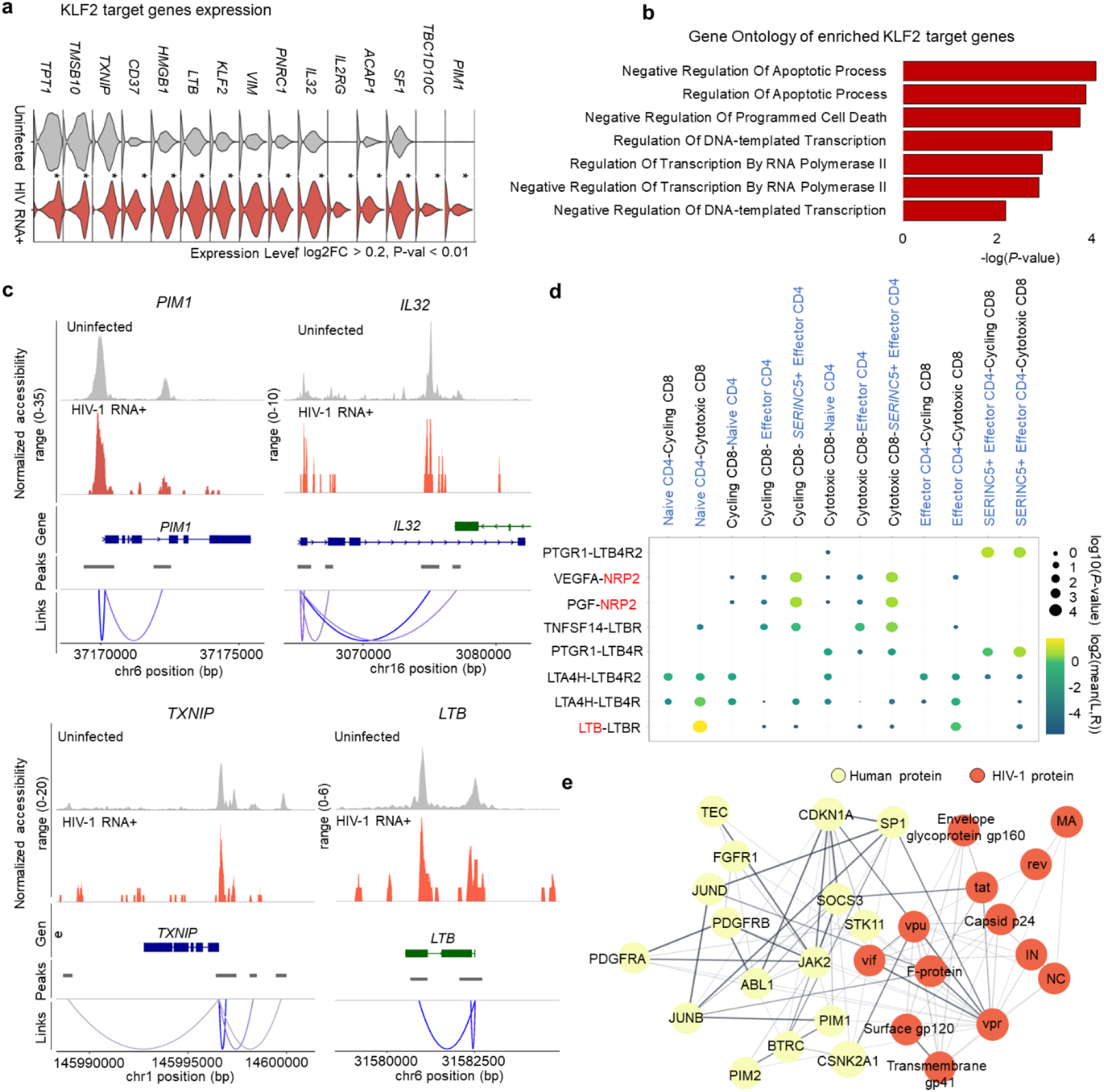
Functional Insights into KLF2 Upregulation in HIV-1 RNA+ CD4 T Cells. (a) Bar plot displays gene ontologies associated with upregulated KLF2 TF target genes in HIV-1 RNA+ cells (*P*-value < 0.01); significance expressed as −log(*P*-value). (b) Violin plot represents expression of upregulated KLF2 TF target genes in uninfected cells and HIV-1 RNA+ CD4 T cells (\**P* < 0.01). (c) Tracks display normalized chromatin accessibility at promoter loci of *PIM1*, *IL32*, *TXNIP*, and *LTB* for HIV-1 RNA+ and uninfected cells, with peak-to-gene links bottom of the coverage plot. (d) Dot plot displays ligand-receptor pairs detected between CD4 and CD8 T cells; red-labeled gene symbols are KLF2 target genes. Circle size indicates *P*-values, while color represents mean average expression levels. (e) Interactions between KLF2 target genes and HIV-1 proteins identified using STRING database. Yellow dots represent human proteins, and red dot represents HIV-1 protein; line thickness indicates interaction strength.

Considering that CD8+ T cells play a critical role in the immune response against HIV-1 infection during chronic infection [47, 48], we further attempted to identify specific ligand– receptor pairs between CD4+ T cells and cytotoxic T lymphocytes to characterize the immune response induced by HIV-1-infected cells [47]. Toward this end, we computed significant ligand–receptor pair interactions between CD8+ and CD4+ T cells using CellPhoneDB (Fig. 4D) [49]. The results indicated a strong interaction between LTB and its receptor (LTBR) in naïve CD4+ T cells, whereas other CD4+ T cell subtypes displayed a weaker interaction with cytotoxic CD8+ T cells. LTB expression was also significantly elevated in HIV-1 RNA+ cells (Supplementary Table 2). Additionally, the NRP2 receptor exhibited heightened levels of interaction between CD4+ and CD8+ T cells, particularly in the effector CD4+ T cell population. *NRP2*, a known target gene of KLF2, is expressed on T cells and modulates immune responses under pathological conditions [50]. Given that KLF2 is involved in immune cell trafficking and adhesion, we used CellChat (v2.1.1) [51] to identify altered ligand–receptor interactions between HIV-1-infected CD4+ T cells and innate immune cells during infection. Both effector and central memory CD4+ T cells exhibited increased MIF and ICAM2 signaling with NK cells, which are associated with KLF2-mediated immune modulation and may enhance immune activation by facilitating immune cell adhesion and migration (Supplementary Fig. 5A) [52, 53]. However, HIV-1-infected CD4+ T cells simultaneously showed reduced CCL5 and CLEC2B signaling, potentially limited monocyte-driven responses and NK cell recruitment [54, 55]. These changes in immune interaction may represent a balance between immune activation and evasion, contributing to viral persistence.

Subsequently, we explored the interactions between the host proteins encoded by KLF2 target genes and HIV-1 proteins. Using the STRING database, we identified 16 KLF2 target genes that encode proteins that directly interact with HIV-1 (Fig. 4E). One of these target genes, *SOCS3* which encodes a protein exhibiting a strong interaction with the Tat protein, pivotal for viral genome replication [56]. Another KLF2 target gene, *PIM1*, which is known as a transactivator of Tat [45], displayed substantial interaction with JUNB and exhibited high gene expression and chromatin accessibility in HIV-1 RNA+ cells (Fig. 4C).

### Th17 cells exhibit increased enrichment and susceptibility to HIV-1 infection

*In vivo* analysis of the states of cells with the HIV-1 provirus integrated into host genome has been a long-standing challenge in HIV research. To address this issue, we defined cells with two or more scATACseq reads that aligned to the HIV-1 genome as HIV-1 DNA+ cells. The sequencing reads predominantly aligned to the 5′- and 3′-long terminal repeat (LTR) regions of the HIV-1 genome (Fig. 5A). We observed clustered HIV-1 DNA+ cells in a distinct region, indicating that these cells share a similar epigenetic profile (Fig. 5B, Supplementary Fig. 6A). DARs were identified in the cluster enriched with HIV-1 DNA+ CD4+ T cells compared to the remaining CD4+ T cells (Fig. 5C). The genes closest to the DARs that showed increased accessibility levels in the HIV-1 DNA+ CD4+ T cell cluster compared to the corresponding levels in the uninfected clusters were associated with the inflammatory and defense responses, which was in line with the activation of the innate immune response observed in HIV-1 RNA+ cells (Fig. 5D).

**Fig. 5.**
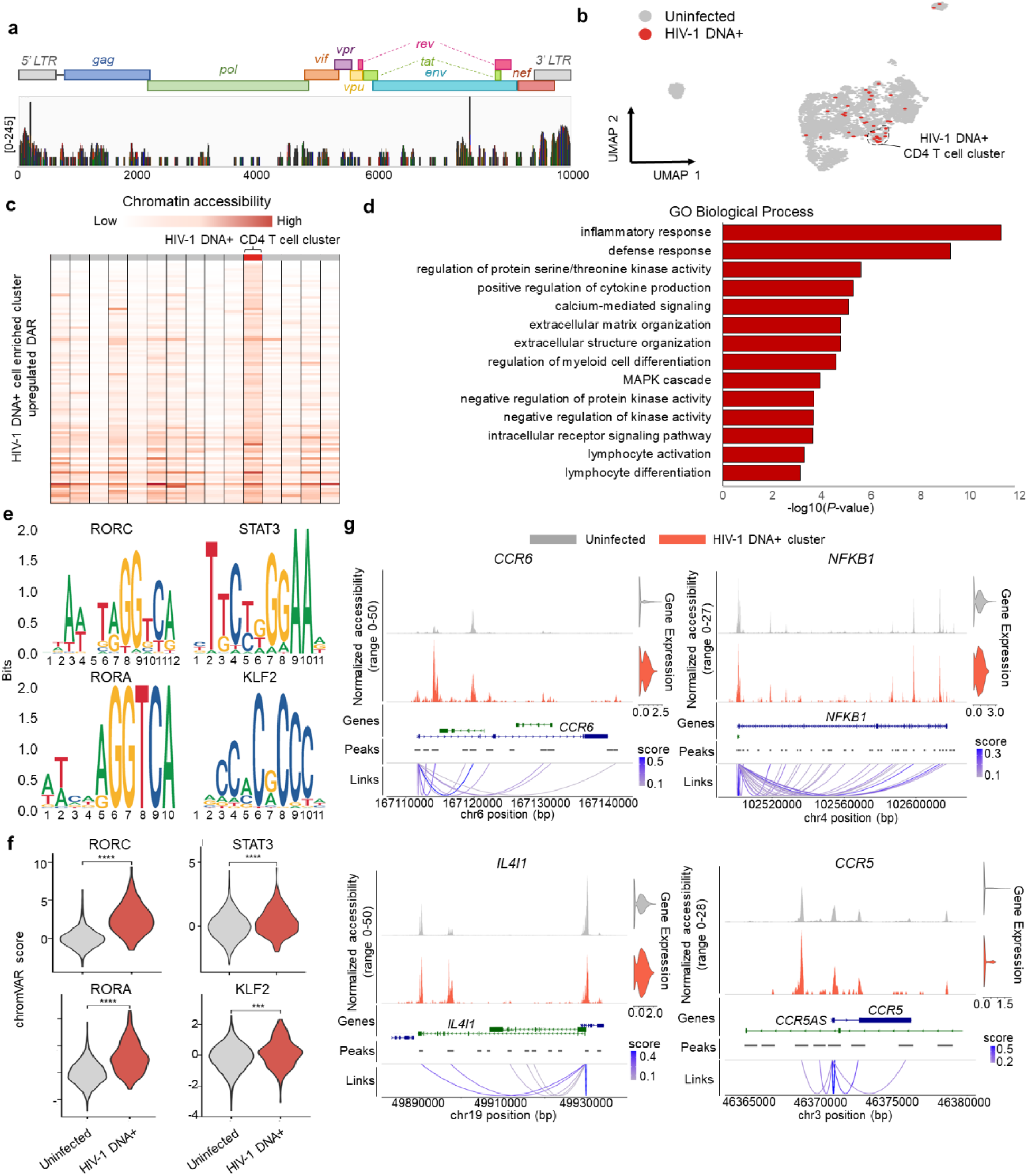
Epigenetic analysis in HIV-1 DNA+ CD4 T cells. (a) Bar plot presents scATAC-seq reads aligned to the HIV-1 genome, showing coverage for individual genomic windows (log-scaled, max value: 245). (b) UMAP of CD4 T cells based on chromatin accessibility identifies 5,535 uninfected cells and 39 HIV-1 DNA+ cells. (c) Heatmap displays chromatin accessibility levels of DARs enriched in HIV-1 DNA+ CD4 T cell cluster. (d) Bar plot represents gene ontologies associated with genes near DARs of HIV-1 DNA+ CD4 T cell cluster; significance expressed as −log(*P*-value). (e) DNA sequences for overrepresented transcription factor binding motifs in HIV-1 DNA+ CD4 T cell cluster compared to the rest of CD4 T cells. (f) Violin plot displays chromVAR motif activity score for enriched motifs. (g) Bar plots on the left show normalized ATAC reads for individual genes in HIV-1 DNA+ CD4 T cells and uninfected cells. Bottom shows peak-to-gene links; loop color strength signifies significance of links between peak and gene promoter. Right violin plot displays gene expression levels.

Notably, we identified distinctly enriched TF-binding motifs in HIV-1 DNA+ CD4+ T cells and HIV-1 RNA+ CD4+ T cells. The binding motifs for key regulators of Th17 cell differentiation (RORC, STAT3, and RORA) were enriched in HIV-1 DNA+ CD4+ T cells (Fig. 5E, F) [57]. These TFs were also reported to regulate viral gene expression by binding to the HIV-1 LTRs [58]. Overall, HIV-1 DNA+ cells mostly represented the Th17 cell type (Supplementary Fig. 6B, C), in contrast to HIV-1 RNA+ CD4+ T cells, which belonged to various subtypes. Additionally, we confirmed significant enrichment of the KLF2 TF-binding motif in HIV-1 DNA+ CD4+ T cells, as found for HIV-1 RNA+ CD4+ T cells. Furthermore, we detected increased gene expression levels and chromatin accessibility of *CCR6*, *IL4I1*, *NFKB1*, and *CCR5* (Fig. 5G).

## DISCUSSION

In response to early HIV-1 infection, various host immune responses are triggered, contingent upon the subtype of CD4+ T cells. Our study utilized scRNA-seq and scATAC-seq datasets to profile PBMCs from early HIV-1-infected patients, revealing specific immune responses and mechanisms in distinct HIV-1-infected CD4+ T cell subsets that contribute to HIV pathogenesis. We further identified key regulators of HIV-1 persistence in CD4+ T cells, offering new targets for complete viral elimination.

Interferons play a crucial role in the initial antiviral response during early viral infection by upregulating interferon-stimulated genes and activating signaling pathways like JAK/STAT, PI3K, and p53-dependent pathways. These pathways facilitate T-cell differentiation and selective proliferation of antiviral T cells, effectively inhibiting viral replication and clearing infected cells [59]. However, our findings indicate that during acute HIV-1 infection, Th2 and Th17 cells exhibit heightened interferon-alpha responses, despite diminished downstream signaling pathways. In parallel, we observed a higher proportion of HIV-1-infected naïve CD4+ T cells. Although their baseline transcriptional activity is low, previous studies have shown that naïve CD4 T cells are susceptible to HIV-1 infection during the early stage of infection, potentially due to dysfunctional antiviral signaling in resting cells [60, 61]. These findings suggest a potential immunological malfunction induced by HIV-1, hindering host antiviral responses by disrupting interferon target signaling, akin to strategies observed in other viruses like Influenza, HCV, and HSV [62].

In CD4+ memory T cells, we observed enriched chromatin accessibility linked to Tat-mediated HIV-1 elongation, concurrent with downregulated interferon-gamma signaling. This underscores HIV-1’s ability to persist in memory CD4+ T cells and evade host antiviral responses early in infection. Through integrated analysis, we identified KLF2 as a pivotal transcription factor within the regulatory network of HIV-1 RNA+ CD4+ T cells. KLF2, known for promoting cellular quiescence and regulating T-cell migration, exhibited sustained expression across all HIV-1-infected CD4+ T-cell subtypes, including activated cells [39]. Upregulated KLF2 target genes associated with HIV-1 replication and TXNIP, a cell proliferation inhibitor, further suggest HIV-1’s ability to persist without triggering transcriptional repressors or proliferation. These findings were further supported by analysis of a publicly available scRNA-seq dataset from HIV-1-infected PBMCs, in which a similar upregulation of *KLF2* and KLF2 target genes was observed in HIV-1-infected CD4+ T cells, reinforcing the reproducibility of our results. A recent study identified FOXP1 and GATA3 as transcriptional regulators of HIV-1 RNA expression using both cell line-based and primary CD4+ T cell models [63]. While these findings highlight transcriptional control in HIV-1 latency, we did not observe notable differential expression of FOXP1 or GATA3 in our dataset (Supplementary Fig. 4B). This discrepancy may reflect differences in infection models, stages, or cellular contexts. Nonetheless, both studies underscore the importance of transcriptional regulation in sustaining HIV-1 persistence.

Using scATAC-seq data, we differentiated HIV-1 DNA+ CD4+ T cells, revealing enriched TF motifs (RORC, STAT3, RORA) indicative of Th17 cell predominance. These cells, unlike HIV-1 RNA+ CD4+ T cells distributed across various subtypes, exhibited heightened susceptibility and integration of HIV DNA, with increased expression of *CCR6* and *CCR5*—critical HIV-1 co-receptors [64, 65]. Moreover, enriched KLF2 motifs in HIV-1 DNA+ cells positively regulate *CCR5* expression, along with increased *IL4I1* expression linked to Th17 cell immunosuppression and *NFKB1* involved in HIV-1 genome transcription initiation [66, 67]. This suggests Th17 cells’ potential role as reservoirs due to their susceptibility, quiescent nature, and stem cell-like characteristics.

Limitations include the inability of our HIV-1 targeted sequencing strategy to distinguish HIV-1 RNA from DNA within single cells, necessitating deeper sequencing to capture transcriptionally silenced proviruses. Our study’s design focused on early infection stages (< 6 months), limiting insights into longitudinal changes in cell populations and gene expression profiles during HIV-1 progression. Given the heterogeneity of HIV-1 infection among PLWH, our findings primarily apply to early infection stages and require validation across different infection phases.

In summary, our study unveils disrupted immune mechanisms in distinct HIV-1 RNA+ CD4+ T cell subsets and underscores KLF2’s role in HIV-1-infected cells. We propose that this dysregulation enables HIV-1 transcription maintenance while attenuating pathogenic immune responses. Furthermore, Th17 cells may serve as reservoirs due to their heightened susceptibility and KLF2-mediated CCR5 expression. Our findings provide insights into the complex transcriptional and epigenetic changes during HIV-1 infection, informing potential immunomodulatory therapies to restore normal immune responses and enhance viral control in PLWH, thereby limiting disease progression.

## METHODS

### Preparation of PBMCs samples from early infection of PLWH

PBMCs were collected from a total of 9 individuals in the early stages of HIV infection (<6 months). Epidemiological and virological characteristics of these participants at baseline, including sex, age, duration from HIV-1 infection, CD4 T cell count, HIV-1 viral load and HIV-1 p24 antigen/antibody screening (Supplementary Table 1).

### Ethics statement

Written informed consent was given by all participants, and the study adhered to the principles of the Declaration of Helsinki and Korea Disease Control and Prevention Agency (IRB No. 2019-07-06-P-A) approved this study.

### Preparation of scRNA-seq libraries

The libraries were prepared using the Chromium Single Cell 3′ Library & Gel Bead Kit v2 (PN-120237) (10x Genomics) following the manufacturer’s instructions. The single-cell suspension was washed twice with 1x PBS containing 0.04% BSA. Cell number and viability were determined using the Bio-Rad TC20 cell counter. The cells were loaded onto the 10x Genomics Chromium Controller to generate gel beads in emulsions (GEMs). Library preparation was performed according to the 10× Genomics Chromium Single Cell 3′ reagent kit (V2 chemistry) instructions. The quality and concentration of the libraries were assessed using the Agilent Bioanalyzer 2100. Finally, the libraries were sequenced on an Illumina NovaSeq.

### Preparation of scATAC-seq libraries

The libraries were prepared using the Chromium Next GEM Single Cell ATAC Library & Gel Bead Kit (PN-1000175) following the manufacturer’s instructions. After isolating the nuclei from PBMCs, the single-nuclei suspension was mixed with Transposition Mix and incubated for 60 minutes at 37°C. Subsequently, the transposed nuclei were combined with the master mix and loaded onto the 10x Genomics Chromium Controller to generate gel beads in emulsions (GEMs). Library construction was performed according to the instructions provided with the Chromium Next GEM Single Cell ATAC Reagent Kits (v1.1). The subsequent steps were performed in the same way as described in ‘Preparation of scRNA-seq libraries’.

### Preparation of single cell multiome libraries

The libraries were prepared using the Chromium Next GEM Single Cell Multiome ATAC + Gene Expression Reagent Kits (PN-1000283) (10x Genomics) following the manufacturer’s instructions. Briefly, nuclei were mixed with Transposition Mix and incubated for 60 minutes at 37°C. Then these were loaded onto Chromium Next GEM Chip J (10x Genomics) and GEMs were generated according to manufacturer’s instructions. After post-GEM cleaned up, barcoded transposed DNA and barcoded full length cDNA fragments were pre-amplified with PCR. ATAC libraries were constructed using the Library Construction Kit (PN-1000190) and full-length pre-amplified cDNA fragments were additionally amplified via PCR. RNA libraries were constructed using the Library Construction Kit B (PN-1000279). Finally, the libraries were sequenced on an Illumina NovaSeq 6000.

### HIV-1 targeted sequencing

After scRNA-seq, remaining scRNA-seq full-length cDNA libraries were subjected to HIV-1 targeted sequencing using the SQK-LSK-109 Ligation Sequencing Kit (Oxford Nanopore Technologies) and three specific primers for HIV-1 (Supplementary Table 3). The TrueSeq Read1 primer was used to preserve the cell barcode information, and the cDNA fragments containing the HIV-1 sequence were amplified by the HIV-1 specific primers. The cDNA fragments with HIV-1 were primarily selected through PCR using biotinylated primers. Subsequently, the HIV-1 specific primers selectively amplified the cDNA fragments through hemi-nested-PCR. These selected cDNA fragments were sequenced on a MinION FLO-MIN106 R9.4.1 flow cell (Oxford Nanopore Technologies). Base calling was performed by using Guppy (V6.1.1), and cells with HIV-1 reads were identified using the cell barcode information.

### Sequence alignment to human and HIV-1 genome

The reads from scRNA-seq and scATAC-seq were aligned to GRCh38 human genome (v.3.0.0) using CellRanger count (10x Genomics, v.6.1.2) and CellRanger ARC (v.2.0.0) with default options, respectively. Feature-barcode matrices were used for downstream analysis. To identify the sequencing reads containing the HIV-1 sequence, the reads from scRNA-seq were aligned to the HIV-1 genome using the STARsolo (v. 2.7.1a). We adjusted the mapping percentage parameter from 50 % to 25% to map the short length of HIV-1 sequence included in the sequencing reads.

### scRNA-seq data processing

For the scRNA-seq datasets, Seurat R package (v.3.2.1) was used for processing the raw count matrices (UMI counts per gene per cell) [68]. Prior to feature filtering, DoubletFinder was used to identify and remove putative doublets[69]. As in previous studies, standard quality control and preprocessing steps were performed for downstream analysis [70, 71]. Cells with fewer than 200 features and greater than 30% mitochondrial gene expression were excluded, and cells expressing fewer than three genes were eliminated. To ensure consistency in quality control and analysis, published scRNA-seq datasets were processed using the same Seurat-based pipeline as described above. Subsequently, the dataset was normalized, highly variable genes were selected, and scaling was performed in Seurat. Batch effects were removed using the ‘RunFastMNN’ function, and principle component analysis (PCA) was performed after integrating the datasets. Cells were clustered based on the computed nearest neighbor map using the ‘FindNeighbors’ function. The clustered cells were visualized on UMAP embeddings. Differentially expressed genes in each cluster were identified using the ‘FindAllMarkers’ function, and the clusters were annotated using the known cell markers. Clusters were merged if the number of differentially expressed genes was less than 10.

### scATAC-seq data processing

For the scATAC-seq datasets, Signac R package (v.1.5.0) was used to process the data.[72] To ensure common features across the datasets, combined peaks were generated. The transcription start site (TSS) enrichment score and nucleosome signal score were calculated for each cell using the ‘TSSEnrichment’ and ‘NucleosomeSignal’ functions in Signac. Cells were retained with a TSS enrichment score > 1 and a nucleosome signal < 2, and further filtered the cells containing < 1000 or > 100,000 ATAC fragments. MACS2 was utilized to identified peaks using each fragment file of datasets.[73] The datasets were merged, and batch effects were removed using Harmony.[74] The merged dataset was subjected to dimensional reduction with LSI using the “RunTFIDF”, “FindTopFeatures” and “RunSVD” functions. The LSI components were used for UMAP projection. The cell type of each cluster was annotated using differentially accessible peaks. To identify significant peaks for each cluster, the ‘FindAllMarks()’ function was employed with the following parameters: min.pct = 0.1, test.use = ‘LR’, only.pos = TRUE. The ‘ClosestFeature()’ function was then used to find the closest genes to each peak within each cluster. Cell clusters were annotated using known cell markers. Subsequently, the scRNA-seq dataset and scATAC dataset from the 10x multiome dataset were integrated using identical cell barcodes to ensure the scATAC-seq clusters’ cell type annotation, leveraging the previously annotated scRNA-seq cell types.

### Gene ontology enrichment analysis and gene set enrichment analysis

To identify enriched biological pathways in HIV-1 RNA+ CD4 T cells, DEGs for each cell type were identified using ‘FindMarkers’ function in Seurat (v.3.2.1). We compared the normalized gene expression data of HIV-1 RNA+ CD4 T cells with uninfected CD4 T cells. All genes that passed quality control were included in the DEG analysis without additional filtering.

For gene ontology enrichment analysis, DEGs were filtered based on |logFC| > 0.25 and adjusted P-value < 0.01. Gene ontology enrichment analysis was performed using the web-based tools of DAVID (version DAVID 6.8; https://david.ncifcrf.gov/). Biological processes (BPs) were ranked based on the sets of DEGs in HIV-1 RNA+ cell, considering BPs with a P-value < 0.01 significant.

For Gene set enrichment analysis (GSEA), the raw count matrix was used, and no additional filtering was applied. GSEA was performed to identify the statistically significant gene sets in the HIV-1 RNA+ cell group within different CD4 T cell subtypes. Hallmark gene sets from Molecular Signatures Database (MSigDB) were used, andonly gene sets containing 10 to 500 genes were selected.

### Gene regulatory network analysis

To construct a gene regulatory network and predict TF activities from a gene expression dataset, SCENIC (R package, https://github.com/aertslab/SCENIC) was utilized as a computational method.[75] Co-expression modules between TFs and target genes were identified using GENIE3. The cisTarget Human motif database was utilized for scoring the regulons. Subsequently, the AUCell method was employed to score single cells and identify enrichment of regulons in the infection group.

### Cell-cell interaction analysis

CellphoneDB was utilized to analyze cell-cell communication between CD4 T cell subtypes and CD8 T cell subtypes.[49] Interaction pairs with a positive mean expression level in the HIV-1 RNA+ group were selected to explore associations among cell types. The interaction strength was visualized using the ‘plot_cpdb’ function in the ktplots R package, represented as a dot plot. To identify dysregulated ligand–receptor signaling, we additionally applied CellChat (v2.1.1) [51]. Differential expression analysis was performed between HIV-1 RNA+ and uninfected CD4+ T cells using the ‘identifyOverExpressedGenes’ function. Upregulated and downregulated interactions were determined based on fold-change values.

### Transcription factor binding motif enrichment and footprinting analysis

The cells were categorized into three groups: HIV-1 RNA+ cell group, HIV-1 DNA+ cell group, and uninfected cell group. Differential accessibility regions (DARs) among the groups were identified using the ‘FindMarkers’ function. DARs with a p-value less than 0.01 between the cell groups were filtered. DARs with a positive log2 fold change value were selected for further analysis to identify enriched motifs. Enriched motifs were identified using the motifmatchr (R package, https://bioconductor.org/packages/motifmatchr), which involved testing for overrepresentation of each DNA sequence motif in the DARs using a hypergeometric test. The computation of GC contents and matching of background sets of peaks for GC content, sequence length, and counts were performed using the ‘FindMotifs’ function in Signac. The per-cell motif activity score was calculated by chromVAR for each motif across the infection conditions.[76] For the identification of a strong enrichment of Tn5 integration events adjacent to the enriched motifs, TF footprinting analysis was conducted using the ‘Footprint’ function in Signac.

### Statistical analysis

The subsection above provides a description of statistical analyses for single cell transcriptomic and epigenetic studies. All descriptive statistical analyses were conducted in R 3.6.0 and higher. Benjamini-Hochberg correction was used for significance.

## ACKNOWLEDGMENTS

We thank all study participants by donating their blood and made this study possible. We are grateful to Jaehyun Seong for preparing this study samples of acute HIV-infected patients. The study samples of acute HIV-infected patients are collected by the Chronic Infectious Disease Cohort Study (Grant Number 4800-4859-304).

## CONTRIBUTORS

Conceptualization: JP, BSC

Data collection: SYC, SSI

Data curation: DL

Formal analysis: DL

Investigation: CHY, SIS, HA

Funding acquisition: JP, BSC

Supervision: JP, BSC

Writing – original draft: DL

Writing – review & editing: JP, BSC, DL

## DECLARATION OF INTERESTS

Authors declare that they have no competing interests.

## FUNDING

This work was supported by the Korea National Institute of Health (Grant Number: 2019-NI-067and 2020-ER5103), the National Research Foundation of Korea (NRF), funded by the Korean government (Grant Number: RS-2024-00335026), and a grant of the Korea-US Collaborative Research Fund(KUCRF), funded by the Ministry of Science and ICT and Ministry of Health & Welfare, Republic of Korea (Grant Number: RS-2024-00466906).

## DATA AVAILABILITY STATEMENT

The raw data was deposited in Korean Nucleotide Archive (KoNA, https://kobic.re.kr/kona)* with the accession ID, KAP230707.

## Supplementary figures

**Fig. S1.**
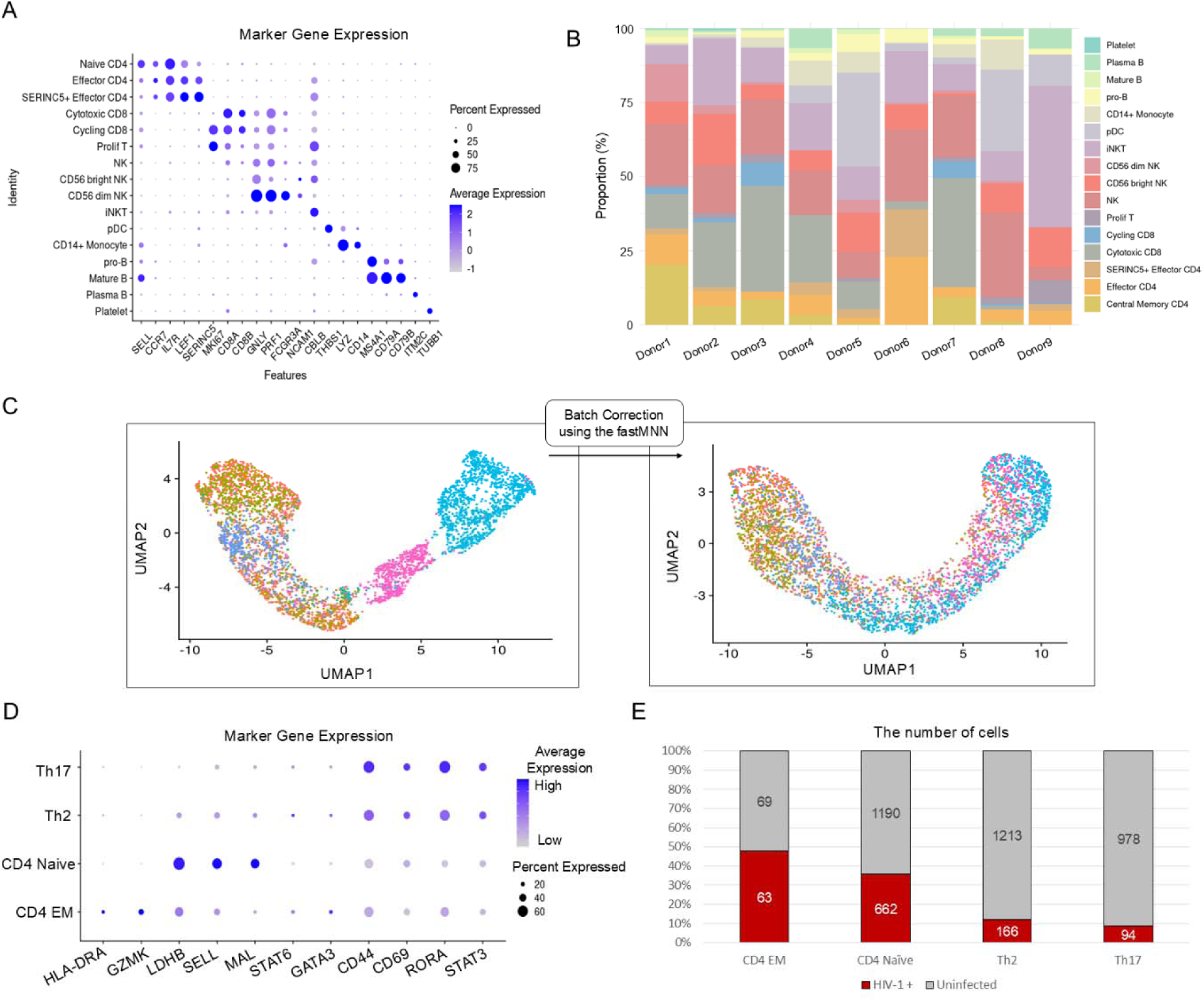
Single-cell transcriptomic analysis identifies the cell types of PBMCs and subclustered CD4 T cells from early HIV-1 infected patients. **(A)** Dot plot illustrates the gene expression of cell type-specific marker genes in all clusters, where the dot size represents the percentage of cells expressing the marker gene, and the color intensity indicates the average expression of the marker gene. **(B)** The proportion of cell types are depicted for each donor. **(C)** UMAP plots of subclustered CD4+ T cells from single-cell RNA-seq data. Each color represents a different donor. The right panel shows the distribution of cells after batch correction using the fastMNN algorithm. **(D)** Dot plot illustrates the gene expression of cell type-specific marker genes in all CD4 subclusters. **(E)** The proportion of HIV-1 RNA+ cells to uninfected cells are depicted for each cell type. The box plot includes the counts of both HIV-1 RNA+ cells and uninfected cells

**Fig. S2.**
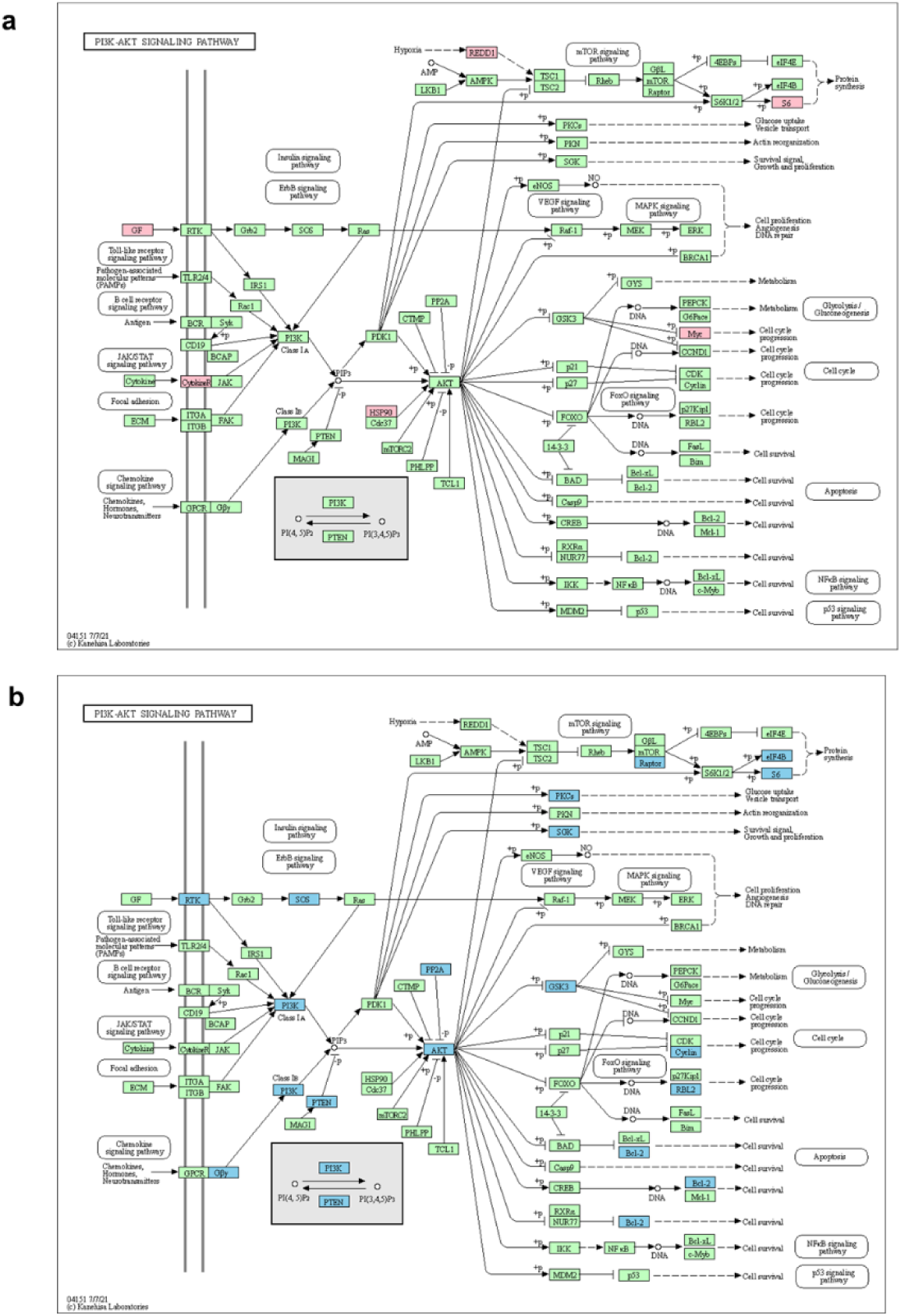
Single-cell transcriptomic analysis identifies the characteristics of HIV-1 RNA+ cells. **(A-B)** Upregulated (red) and downregulated (blue) genes are shown on the map of KEGG PI3K-AKT signaling pathway.

**Fig. S3.**
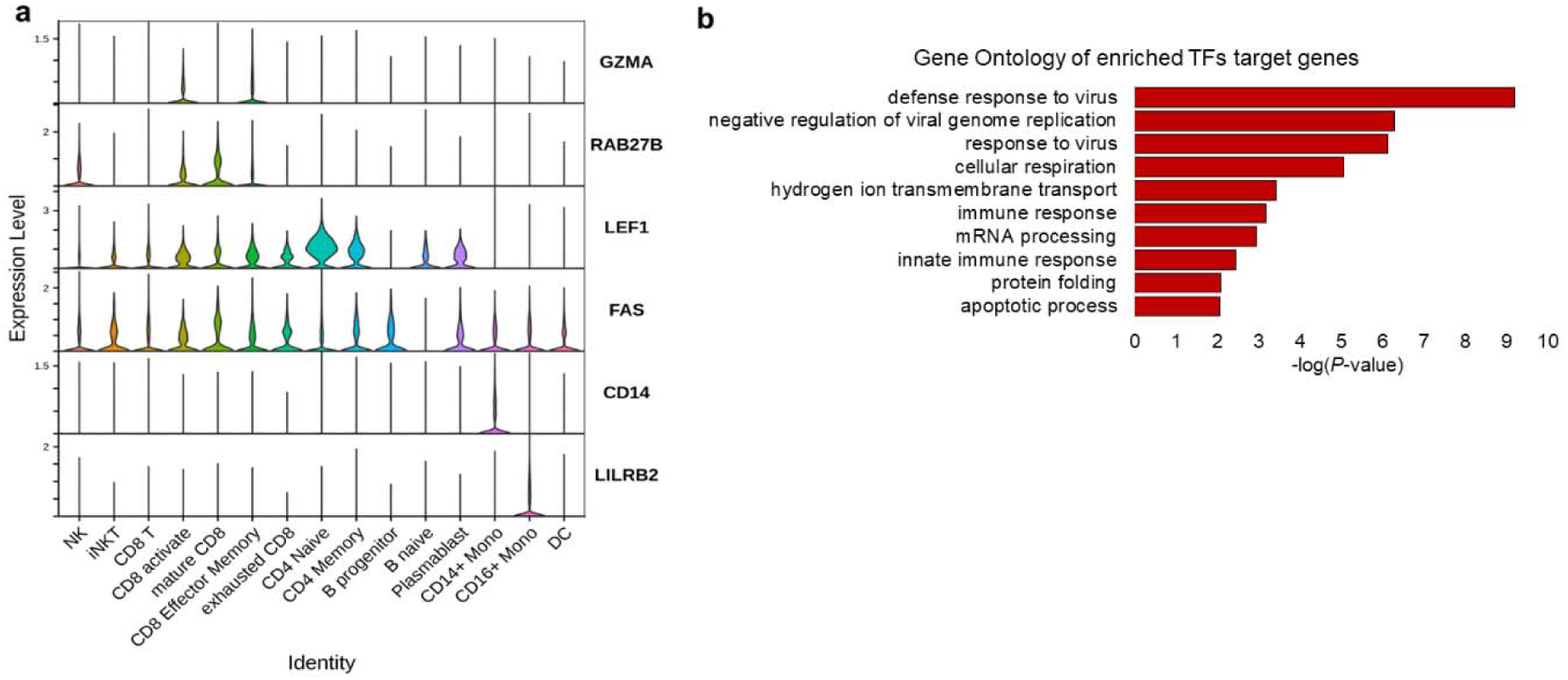
Single-cell multi-omics analysis identifies epigenetic characteristics of scATAC-seq dataset and upregulated transcription factors in HIV-1 RNA+ cells. **(A)** Violin plot displays the gene expression of marker genes in scATAC-seq dataset. **(B)** The bar plot represents the gene ontologies associated with the target genes of the upregulated TFs in HIV-1 RNA+ cells (*P*-value < 0.01). The significance of these enrichments is expressed as −log(*P*-value).

**Fig. S4.**
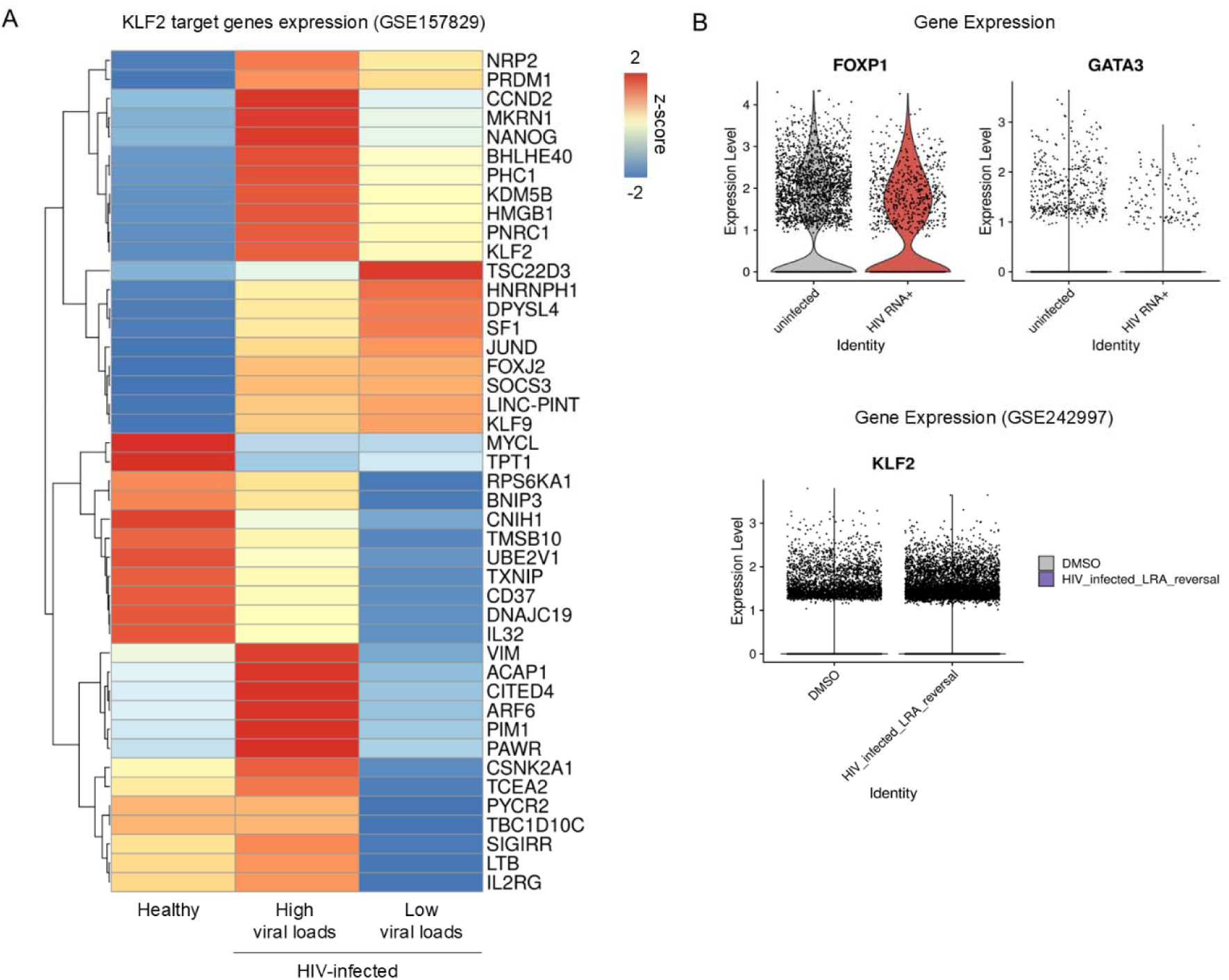
KLF2 target gene expression in HIV-1-infected CD4+ T cells. **(A)** Heatmap showing the expression (Z-score normalized) of KLF2 target genes across healthy controls and HIV-1-infected individuals with high and low viral loads. Gene expression data were obtained from a published dataset (GSE157829) **(B)** Violin plots showing the expression levels of *FOXP1* and *GATA3* in HIV-1 RNA+ versus uninfected CD4+ T cells (top, current study). Bottom panel shows expression levels of *KLF2* in published dataset (GSE242997), comparing HIV-infected cells treated with latency-reversing agents (LRA) and DMSO controls.

**Fig. S5.**
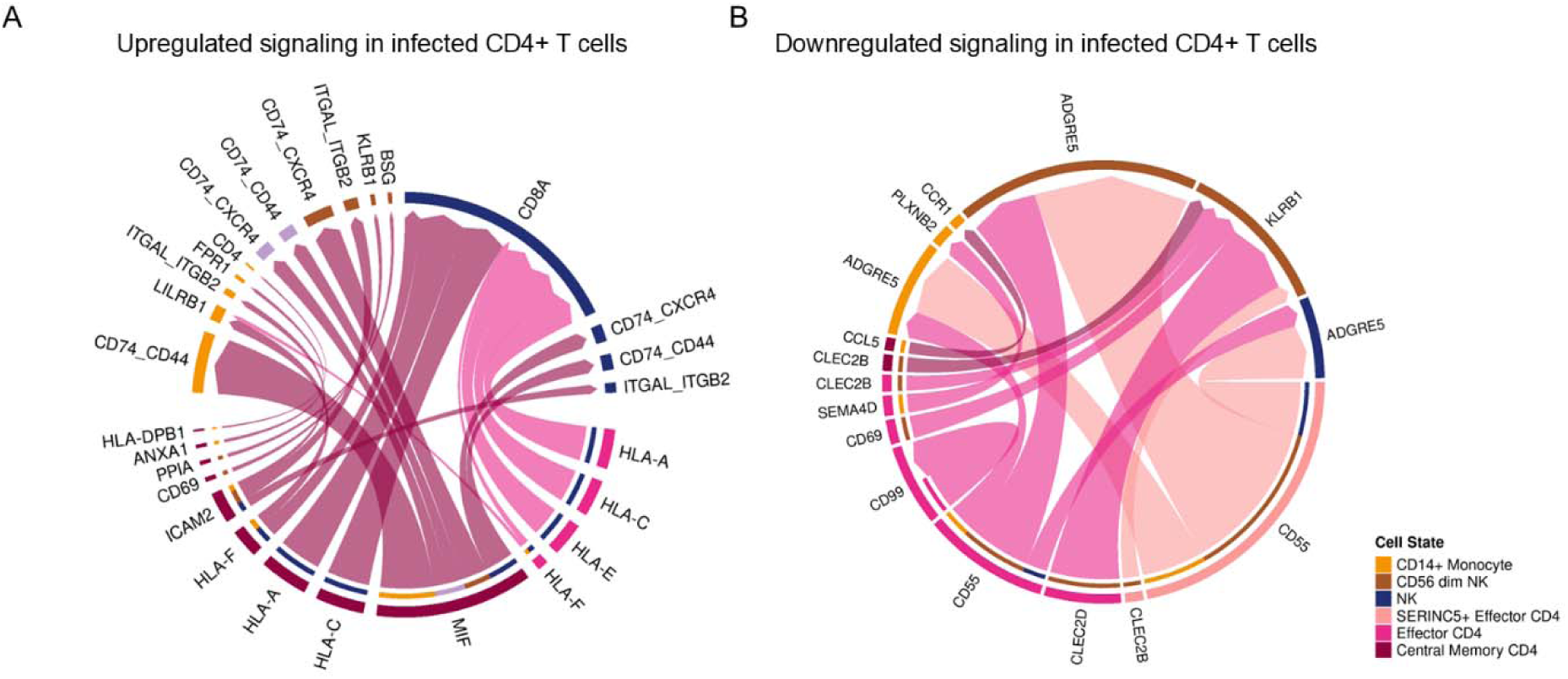
Cell-cell interaction in HIV-1-infected CD4+ T cells. (A) Circos plot show cell–cell interactions that are upregulated in HIV-1-infected CD4+ T cells. Line thickness represents interaction strength, and arcs are colored by cell types. (B) Circos plot shows downregulated interactions in HIV-1 infected CD4+ T cells.

**Fig. S6.**
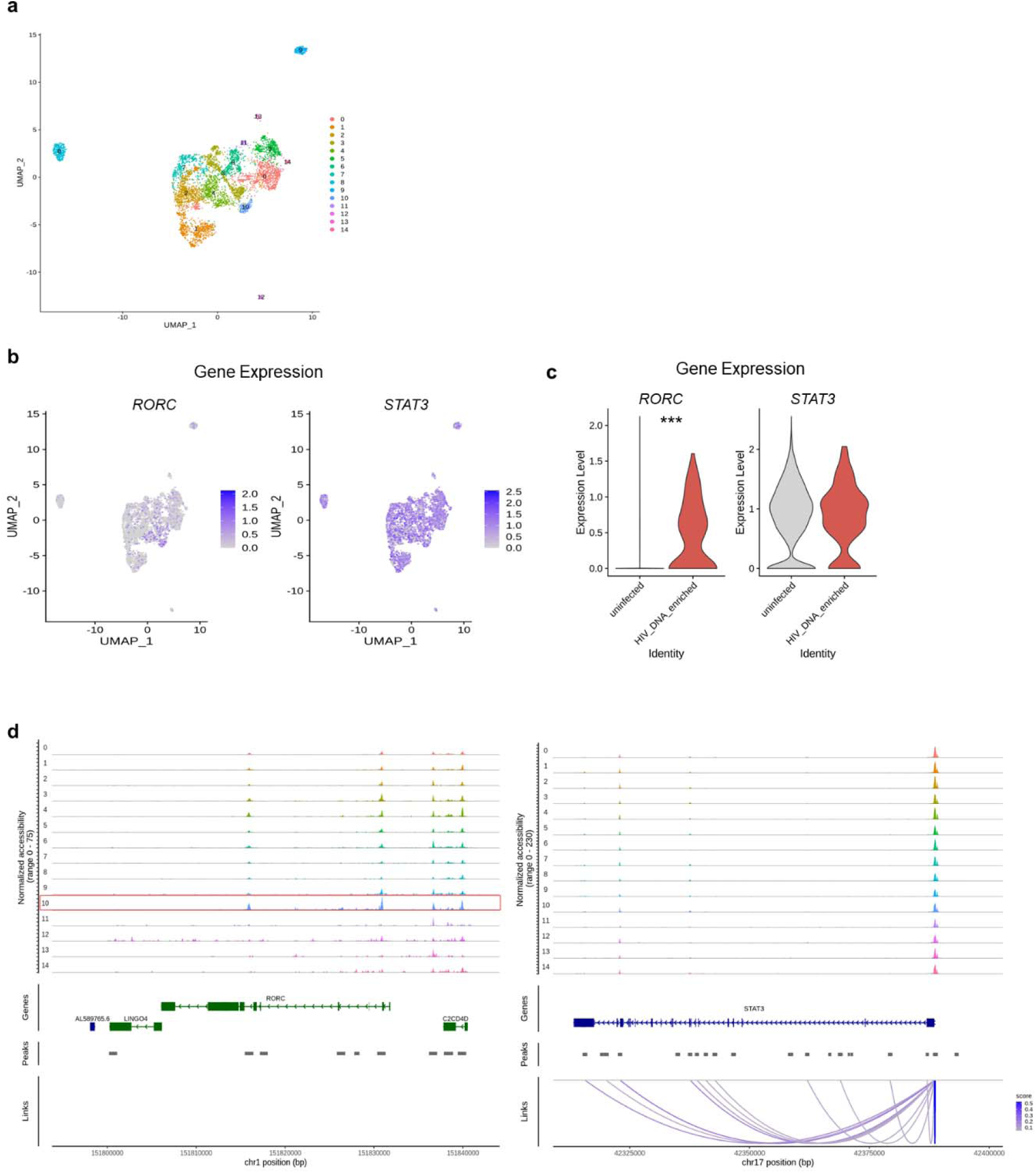
Single-cell epigenetic analysis identifies HIV-1 DNA+ cells are mostly represented by Th17 cell type. **(A)** The ATAC UMAP plot displays the distribution of 5,608 CD4 T cells from early infected patients. **(B-C)** The Feature Plot and violin plot indicate gene expression (*RORC*, *STAT3*) in CD4 T cells. \**P* < 0.01 **(D)** The tracks display the normalized chromatin accessibility levels at the promoter regions of *RORC* and *STAT3* in all cluster of CD4 T cells.

## Supplementary Tables

**Table S1.**
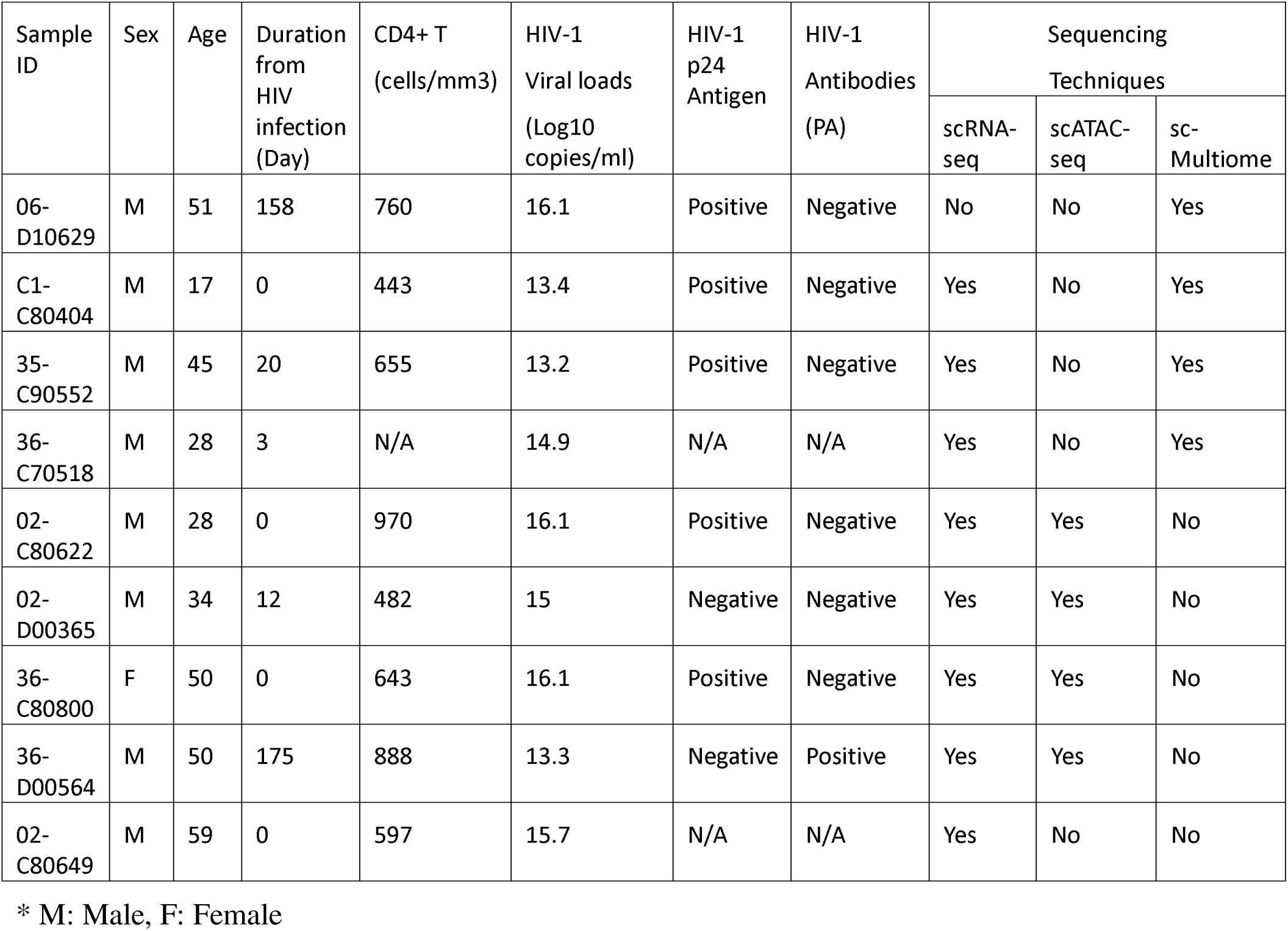
Epidemiological and Virological Characteristics, and Sequencing Data Information for 9 Acute HIV-infected Patients at Baseline.

**Table S2.**
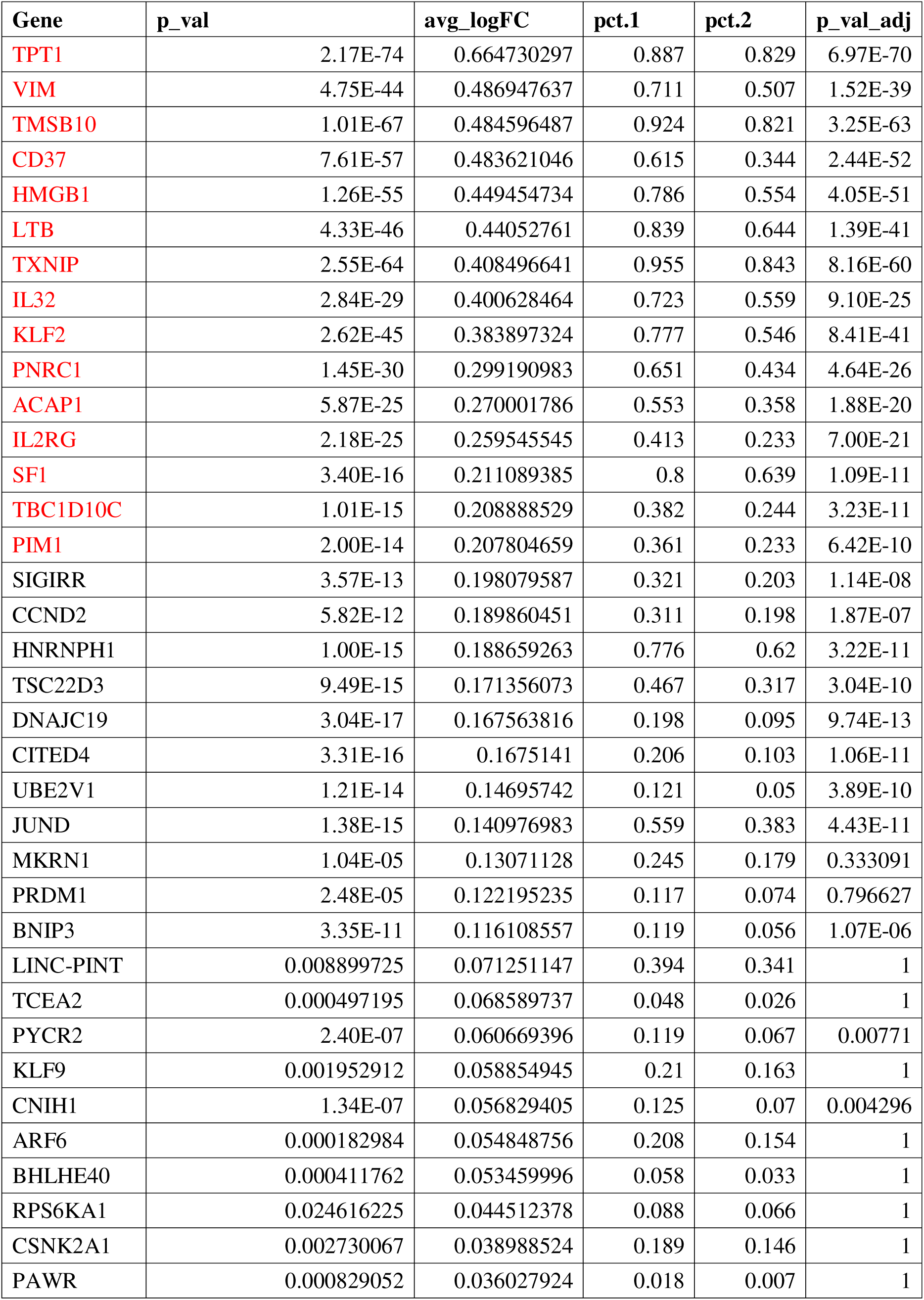
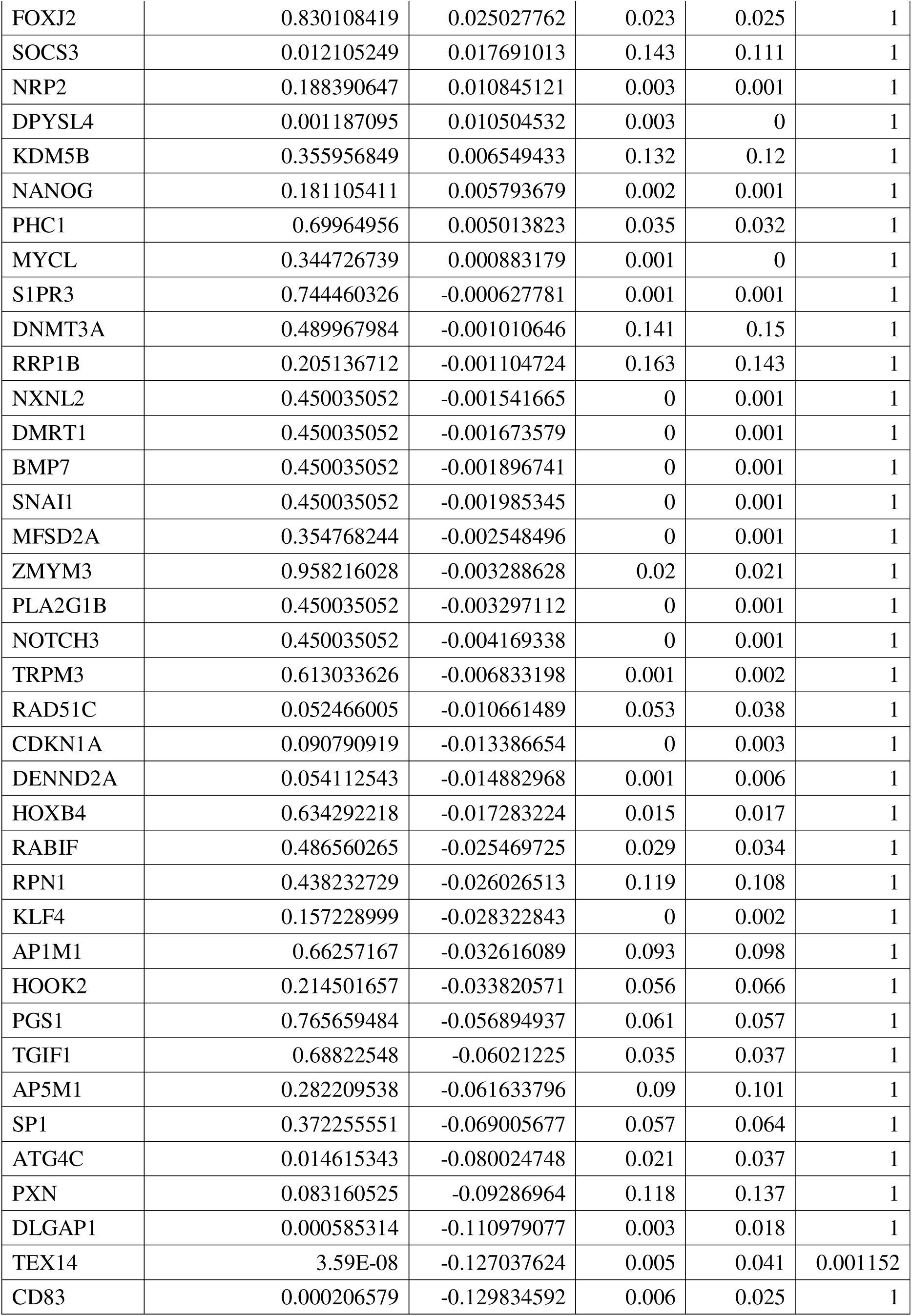
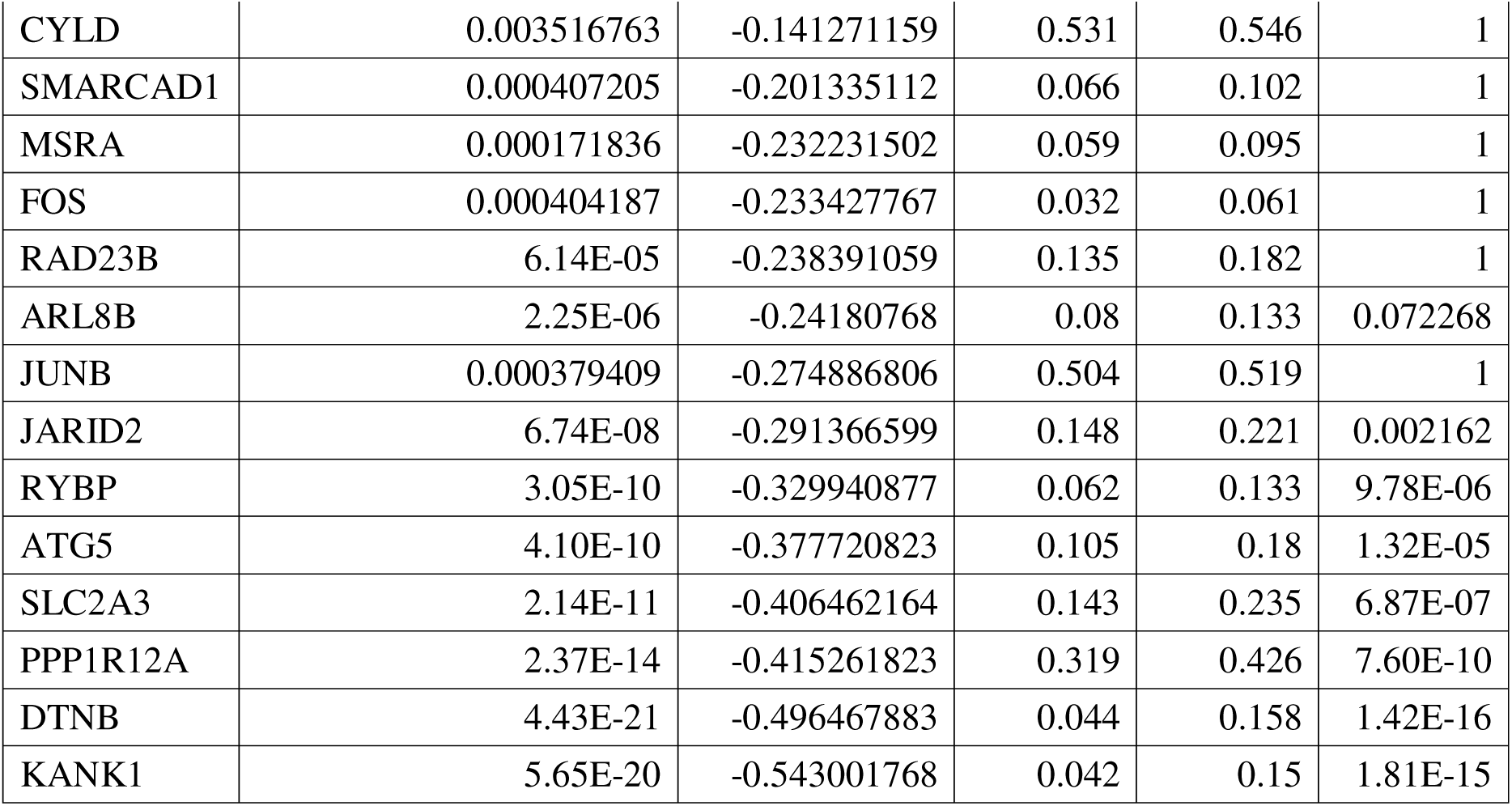
Differentially expressed genes overlapped with KLF2 target gene in CD4 T cell. (HIV-1 RNA+ vs. Uninfected)

**Table S3.**
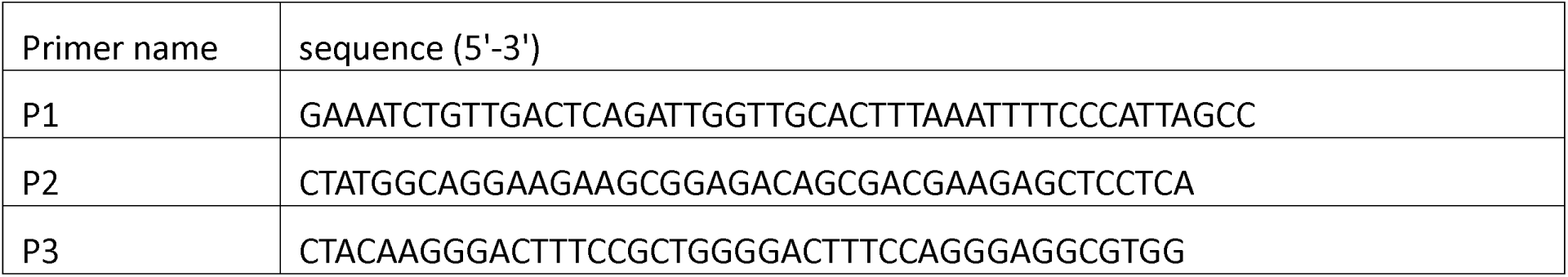
Primers used for HIV-1 targeted sequencing.

## Notes

### Competing Interest Statement

The authors have declared no competing interest.

### Summary of Updates

Section on "The single cell transcriptional landscape of HIV-1 infected cells in early stage of infection", "KLF2 is an essential regulator in the function of HIV-1-infected CD4+ T cells", Discussion and Methods are updated; Funding updated; Acknowledgments updated; Supplemental files updated.

## REFERENCES

1. Douek, D.C., L.J. Picker, and R.A. Koup, T cell dynamics in HIV-1 infection. Annual Review of Immunology, 2003. 21: p. 265–304.

2. Pantaleo, G., C. Graziosi, and A.S. Fauci, The role of lymphoid organs in the pathogenesis of HIV infection. Semin Immunol, 1993. 5(3): p. 157–63.

3. Permanyer, M., et al., Antiretroviral agents effectively block HIV replication after cell-to-cell transfer. J Virol, 2012. 86(16): p. 8773–80.

4. Peterson, T.E. and J.V. Baker, Assessing inflammation and its role in comorbidities among persons living with HIV. Current Opinion in Infectious Diseases, 2019. 32(1): p. 8–15.

5. Zicari, S., et al., Immune Activation, Inflammation, and Non-AIDS Co-Morbidities in HIV-Infected Patients under Long-Term ART. Viruses-Basel, 2019. 11(3).

6. Battistini, A. and M. Sgarbanti, HIV-1 Latency: An Update of Molecular Mechanisms and Therapeutic Strategies. Viruses-Basel, 2014. 6(4): p. 1715–1758.

7. Fiebig, E.W., et al., Dynamics of HIV viremia and antibody seroconversion in plasma donors: implications for diagnosis and staging of primary HIV infection. Aids, 2003. 17(13): p. 1871–1879.

8. Verdikt, R., O. Hernalsteens, and C. Van Lint, Epigenetic Mechanisms of HIV-1 Persistence. Vaccines, 2021. 9(5).

9. Lange, U.C., et al., Epigenetic crosstalk in chronic infection with HIV-1. Seminars in Immunopathology, 2020. 42(2): p. 187–200.

10. Park, J., et al., Genome-wide analysis of histone modifications in latently HIV-1 infected T cells. AIDS, 2014. 28(12): p. 1719–28.

11. Parker, E., et al., Gene dysregulation in acute HIV-1 infection - early transcriptomic analysis reveals the crucial biological functions affected. Frontiers in Cellular and Infection Microbiology, 2023. 13.

12. Deeks, S.G., et al., HIV infection. Nat Rev Dis Primers, 2015. 1: p. 15035.

13. Yoon, B., et al., Single-cell lineage tracing approaches to track kidney cell development and maintenance. Kidney Int, 2024. 105(6): p. 1186–1199.

14. Bae, S.G., et al., Single-cell transcriptome analysis of cavernous tissues reveals the key roles of pericytes in diabetic erectile dysfunction. Elife, 2024. 12.

15. Eun, M., et al., Chromatin accessibility analysis and architectural profiling of human kidneys reveal key cell types and a regulator of diabetic kidney disease. Kidney Int, 2024. 105(1): p. 150–164.

16. Kim, J. and J. Park, Single-cell transcriptomics: a novel precision medicine technique in nephrology. Korean J Intern Med, 2021. 36(3): p. 479–490.

17. Luo, H., et al., Pan-cancer single-cell analysis reveals the heterogeneity and plasticity of cancer-associated fibroblasts in the tumor microenvironment. Nat Commun, 2022. 13(1): p. 6619.

18. Heumos, L., et al., Best practices for single-cell analysis across modalities. Nat Rev Genet, 2023. 24(8): p. 550–572.

19. Lee, D., et al., Transcriptome Analysis Identifies Altered Biological Processes and Novel Markers in Human Immunodeficiency Virus-1 Long-Term Non-Progressors. Infect Chemother, 2021. 53(3):p. 489–502.

20. Bradley, T., et al., Single-Cell Analysis of Quiescent HIV Infection Reveals Host Transcriptional Profiles that Regulate Proviral Latency. Cell Reports, 2018. 25(1): p. 107-+.

21. Bruner, K.M., et al., A quantitative approach for measuring the reservoir of latent HIV-1 proviruses. Nature, 2019. 566(7742): p. 120-+.

22. Kazer, S.W., et al., Integrated single-cell analysis of multicellular immune dynamics during hyperacute HIV-1 infection. Nature Medicine, 2020. 26(4): p. 511-+.

23. Sun, W.W., et al., Phenotypic signatures of immune selection in HIV-1 reservoir cells. Nature, 2023.

24. Wei, Y., et al., Single-cell epigenetic, transcriptional, and protein profiling of latent and active HIV-1 reservoir revealed that IKZF3 promotes HIV-1 persistence. Immunity, 2023. 56(11): p. 2584–2601 e7.

25. Ho, Y.C., et al., Replication-Competent Noninduced Proviruses in the Latent Reservoir Increase Barrier to HIV-1 Cure. Cell, 2013. 155(3): p. 540–551.

26. Liu, R.X., et al., Single-cell transcriptional landscapes reveal HIV-1-driven aberrant host gene transcription as a potential therapeutic target. Science Translational Medicine, 2020. 12(543).

27. Golumbeanu, M., et al., Single-Cell RNA-Seq Reveals Transcriptional Heterogeneity in Latent and Reactivated HIV-Infected Cells. Cell Reports, 2018. 23(4): p. 942–950.

28. Collora, J.A., et al., Single-cell multiomics reveals persistence of HIV-1 in expanded cytotoxic T cell clones. Immunity, 2022. 55(6): p. 1013–1031 e7.

29. Wu, V.H., et al., Profound phenotypic and epigenetic heterogeneity of the HIV-1-infected CD4(+) T cell reservoir. Nat Immunol, 2023. 24(2): p. 359–370.

30. Geretz, A., et al., Single-cell transcriptomics identifies prothymosin alpha restriction of HIV-1 in vivo. Sci Transl Med, 2023. 15(707): p. eadg0873.

31. Lu, J., et al., The IFITM proteins inhibit HIV-1 infection. J Virol, 2011. 85(5): p. 2126–37.

32. Horsburgh, B.A., et al., Cellular Activation, Differentiation, and Proliferation Influence the Dynamics of Genetically Intact Proviruses Over Time. Journal of Infectious Diseases, 2022. 225(7): p. 1168–1178.

33. Apcher, G.S., et al., Human immunodeficiency virus-1 Tat protein interacts with distinct proteasomal α and β subunits. Febs Letters, 2003. 553(1-2): p. 200–204.

34. Wang, H., et al., A computational study of Tat-CDK9-Cyclin binding dynamics and its implication in transcription-dependent HIV latency. Phys Chem Chem Phys, 2020. 22(44): p. 25474–25482.

35. Sreeram, S., et al., The potential role of HIV-1 latency in promoting neuroinflammation and HIV-1-associated neurocognitive disorder. Trends Immunol, 2022. 43(8): p. 630–639.

36. Chen, J., et al., Transcriptome analysis of CD4(+) T cells from HIV-infected individuals receiving ART with LLV revealed novel transcription factors regulating HIV-1 promoter activity. Virol Sin, 2023. 38(3): p. 398–408.

37. Uchil, P.D., et al., TRIM E3 ligases interfere with early and late stages of the retroviral life cycle. PLoS Pathog, 2008. 4(2): p. e16.

38. Masenga, S.K., et al., HIV-Host Cell Interactions. Cells, 2023. 12(10).

39. Kuo, C.T., M.L. Veselits, and J.M. Leiden, LKLF: A transcriptional regulator of single-positive T cell quiescence and survival. Science, 1997. 277(5334): p. 1986–1990.

40. Takada, K., et al., Kruppel-Like Factor 2 Is Required for Trafficking but Not Quiescence in Postactivated T Cells. The Journal of Immunology, 2011. 186(2): p. 775–783.

41. Pedro, K.D., et al., A functional screen identifies transcriptional networks that regulate HIV-1 and HIV-2. Proceedings of the National Academy of Sciences of the United States of America, 2021. 118(11).

42. Hart, G.T., K.A. Hogquist, and S.C. Jameson, Kruppel-like factors in lymphocyte biology. J Immunol, 2012. 188(2): p. 521–6.

43. Wang, S., et al., An atlas of immune cell exhaustion in HIV-infected individuals revealed by single-cell transcriptomics. Emerg Microbes Infect, 2020. 9(1): p. 2333–2347.

44. Duverger, A., et al., Kinase control of latent HIV-1 infection: PIM-1 kinase as a major contributor to HIV-1 reactivation. J Virol, 2014. 88(1): p. 364–76.

45. Didichenko, S.A., et al., IL-3 induces a Pim1-dependent antiapoptotic pathway in primary human basophils. Blood, 2008. 112(10): p. 3949–58.

46. Muri, J., H. Thut, and M. Kopf, The thioredoxin-1 inhibitor Txnip restrains effector T-cell and germinal center B-cell expansion. Eur J Immunol, 2021. 51(1): p. 115–124.

47. Gulzar, N. and K.F.T. Copeland, CD8+T-cells: Function and response to HIV infection. Current Hiv Research, 2004. 2(1): p. 23–37.

48. Mudd, J.C. and M.M. Lederman, CD8 T cell persistence in treated HIV infection. Curr Opin HIV AIDS, 2014. 9(5): p. 500–5.

49. Efremova, M., et al., CellPhoneDB: inferring cell-cell communication from combined expression of multi-subunit ligand-receptor complexes. Nature Protocols, 2020. 15(4): p. 1484–1506.

50. Roy, S., et al., Multifaceted Role of Neuropilins in the immune System: Potential Targets for immunotherapy. Frontiers in Immunology, 2017. 8.

51. Jin, S., M.V. Plikus, and Q. Nie, CellChat for systematic analysis of cell-cell communication from single-cell transcriptomics. Nat Protoc, 2025. 20(1): p. 180–219.

52. Pestal, K., L.C. Slayden, and G.M. Barton, Kruppel-like Factor (KLF) family members control expression of genes required for serous cavity and alveolar macrophage identities. bioRxiv, 2024.

53. SenBanerjee, S., et al., KLF2 is a novel transcriptional regulator of endothelial proinflammatory activation. Journal of Experimental Medicine, 2004. 199(10): p. 1305–1315.

54. Gouwy, M., et al., CXCR4 and CCR5 ligands cooperate in monocyte and lymphocyte migration and in inhibition of dual-tropic (R5/X4) HIV-1 infection. Eur J Immunol, 2011. 41(4): p. 963–73.

55. Welte, S., et al., Mutual activation of natural killer cells and monocytes mediated by NKp80-AICL interaction. Nat Immunol, 2006. 7(12): p. 1334–42.

56. Akhtar, L.N., et al., Suppressor of cytokine signaling 3 inhibits antiviral IFN-beta signaling to enhance HIV-1 replication in macrophages. J Immunol, 2010. 185(4): p. 2393–404.

57. Castro, G., et al., RORgammat and RORalpha signature genes in human Th17 cells. PLoS One, 2017. 12(8): p. e0181868.

58. Wiche Salinas, T.R., et al., Th17 cell master transcription factor RORC2 regulates HIV-1 gene expression and viral outgrowth. Proc Natl Acad Sci U S A, 2021. 118(48).

59. Satarker, S., et al., JAK-STAT Pathway Inhibition and their Implications in COVID-19 Therapy. Postgraduate Medicine, 2021. 133(5): p. 489–507.

60. Douek, D.C., et al., HIV preferentially infects HIV-specific CD4+ T cells. Nature, 2002. 417(6884):p. 95–8.

61. Jiao, Y., et al., Higher HIV DNA in CD4+ naive T-cells during acute HIV-1 infection in rapid progressors. Viral Immunol, 2014. 27(6): p. 316–8.

62. Katze, M.G., Y. He, and M. Gale, Jr., Viruses and interferon: a fight for supremacy. Nat Rev Immunol, 2002. 2(9): p. 675–87.

63. Ashokkumar, M., et al., Integrated Single-cell Multiomic Analysis of HIV Latency Reversal Reveals Novel Regulators of Viral Reactivation. Genomics Proteomics Bioinformatics, 2024. 22(1).

64. Richardson, M.W., et al., Kruppel-like factor 2 modulates CCR5 expression and susceptibility to HIV-1 infection. J Immunol, 2012. 189(8): p. 3815–21.

65. Hutter, G., et al., Long-term control of HIV by CCR5 Delta32/Delta32 stem-cell transplantation. N Engl J Med, 2009. 360(7): p. 692–8.

66. Cousin, C., et al., The immunosuppressive enzyme IL4I1 promotes FoxP3(+) regulatory T lymphocyte differentiation. Eur J Immunol, 2015. 45(6): p. 1772–82.

67. Coiras, M., et al., Basal shuttle of NF-kappaB/I kappaB alpha in resting T lymphocytes regulates HIV-1 LTR dependent expression. Retrovirology, 2007. 4: p. 56.

68. Stuart, T., et al., Comprehensive Integration of Single-Cell Data. Cell, 2019. 177(7): p. 1888–1902 e21.

69. McGinnis, C.S., L.M. Murrow, and Z.J. Gartner, DoubletFinder: Doublet Detection in Single-Cell RNA Sequencing Data Using Artificial Nearest Neighbors. Cell Syst, 2019. 8(4): p. 329–337 e4.

70. Kim, G.D., C. Lim, and J. Park, A practical handbook on single-cell RNA sequencing data quality control and downstream analysis. Mol Cells, 2024. 47(9): p. 100103.

71. Kim, G.D., et al., Cell Type- and Age-Specific Expression of lncRNAs across Kidney Cell Types. J Am Soc Nephrol, 2024. 35(7): p. 870–885.

72. Stuart, T., et al., Single-cell chromatin state analysis with Signac. Nat Methods, 2021. 18(11): p. 1333–1341.

73. Zhang, Y., et al., Model-based analysis of ChIP-Seq (MACS). Genome Biol, 2008. 9(9): p. R137.

74. Korsunsky, I., et al., Fast, sensitive and accurate integration of single-cell data with Harmony. Nat Methods, 2019. 16(12): p. 1289–1296.

75. Aibar, S., et al., SCENIC: single-cell regulatory network inference and clustering. Nat Methods, 2017. 14(11): p. 1083–1086.

76. Schep, A.N., et al., chromVAR: inferring transcription-factor-associated accessibility from single-cell epigenomic data. Nat Methods, 2017. 14(10): p. 975–978.

