## Supplemental Files for "Multi-Omics Single-Cell Analysis Reveals Key Regulators of HIV-1 Persistence and Aberrant Host Immune Responses in Early Infection"

### **Supplementary figure**

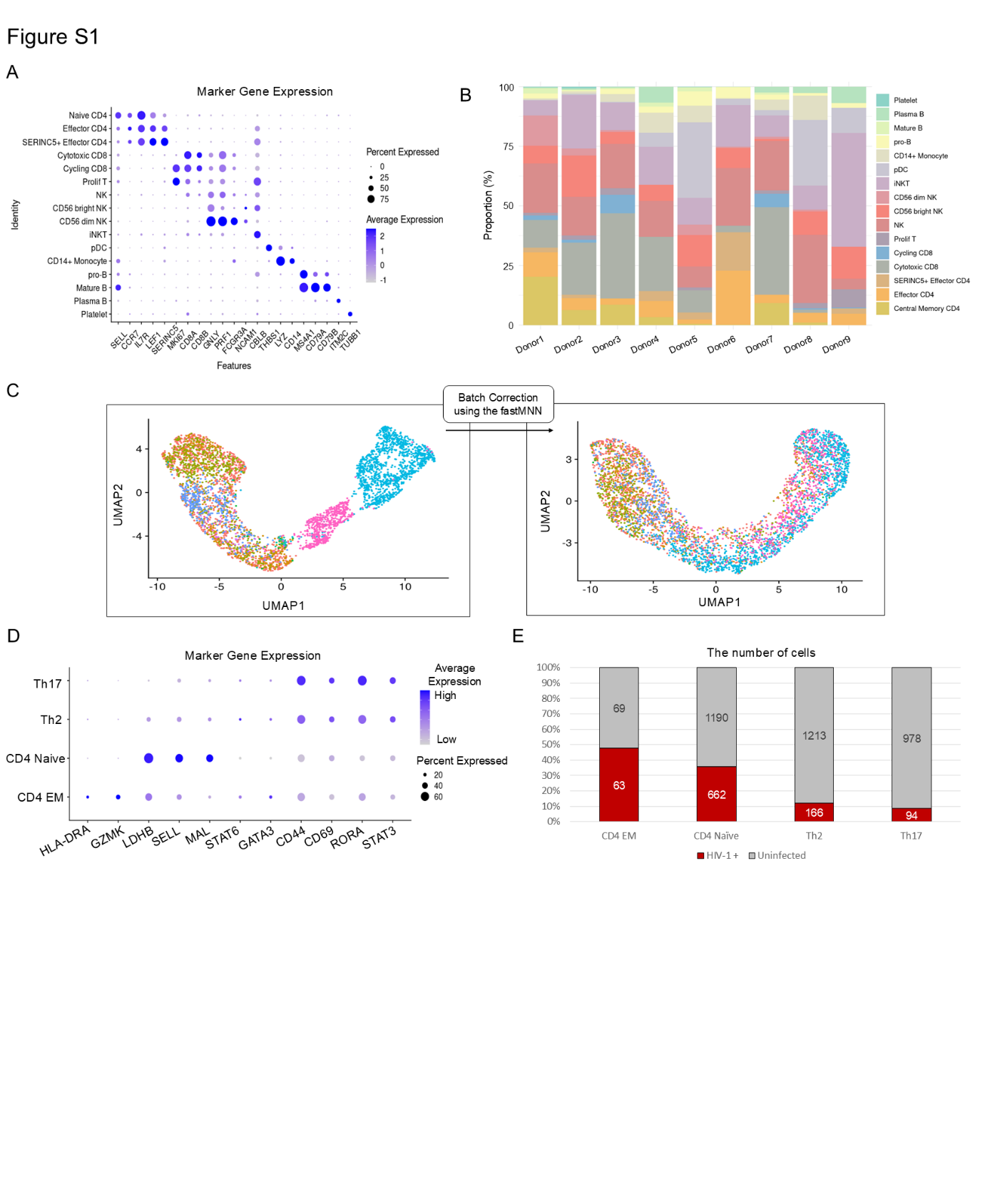

**Fig. S1**. **Single-cell transcriptomic analysis identifies the cell types of PBMCs and subclustered CD4 T cells from early HIV-1 infected patients.**

**(A)** Dot plot illustrates the gene expression of cell type-specific marker genes in all clusters, where the dot size represents the percentage of cells expressing the marker gene, and the color intensity indicates the average expression of the marker gene.

**(B)** The proportion of cell types are depicted for each donor.

**(C)** UMAP plots of subclustered CD4+ T cells from single-cell RNA-seq data. Each color represents a different donor. The right panel shows the distribution of cells after batch correction using the fastMNN algorithm.

**(D)** Dot plot illustrates the gene expression of cell type-specific marker genes in all CD4 subclusters.

**(E)** The proportion of HIV-1 RNA+ cells to uninfected cells are depicted for each cell type. The box plot includes the counts of both HIV-1 RNA+ cells and uninfected cells

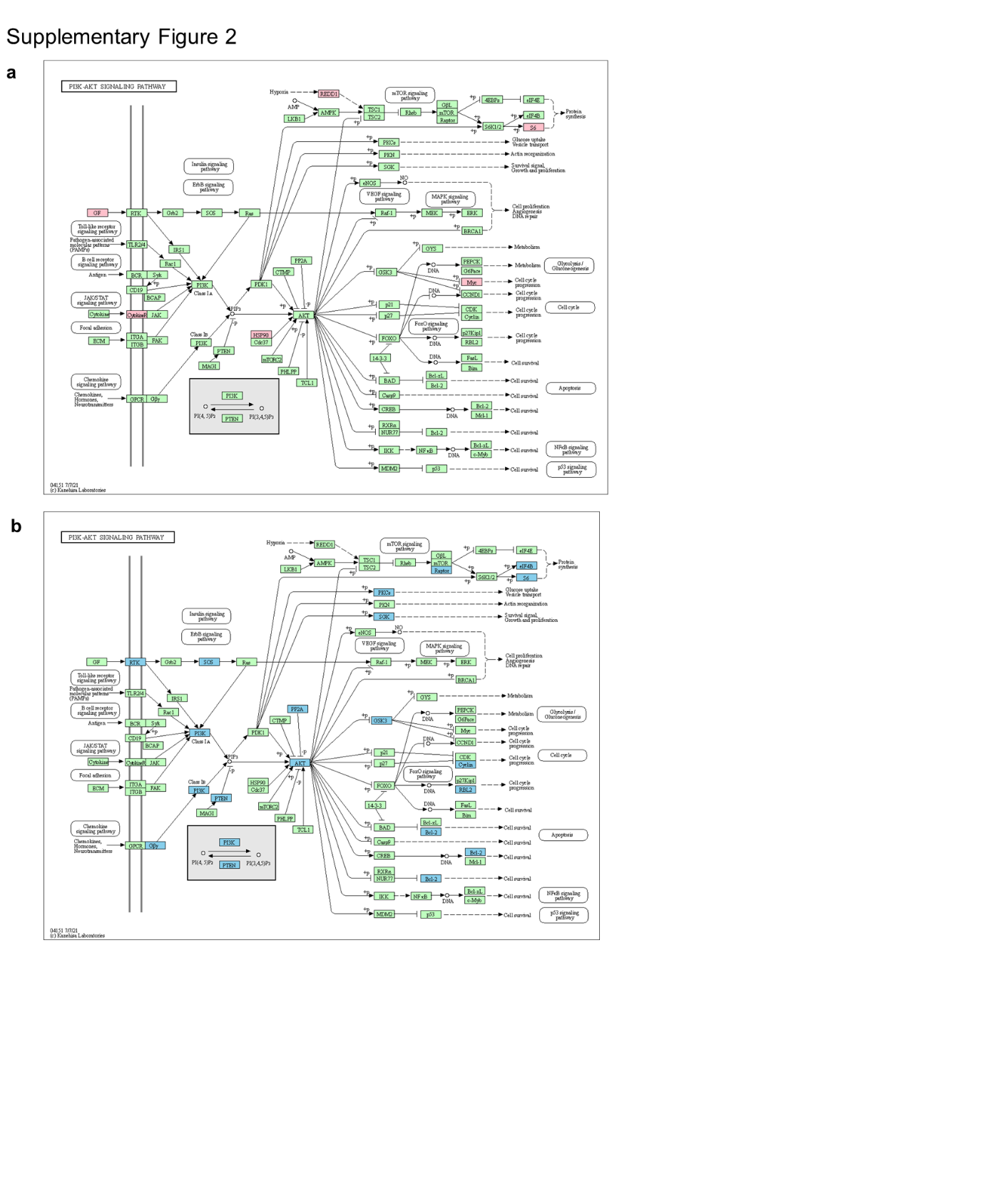
**Fig. S2**. **Single-cell transcriptomic analysis identifies the characteristics of HIV-1 RNA+ cells.**

**(A-B)** Upregulated (red) and downregulated (blue) genes are shown on the map of KEGG PI3K-AKT signaling pathway.

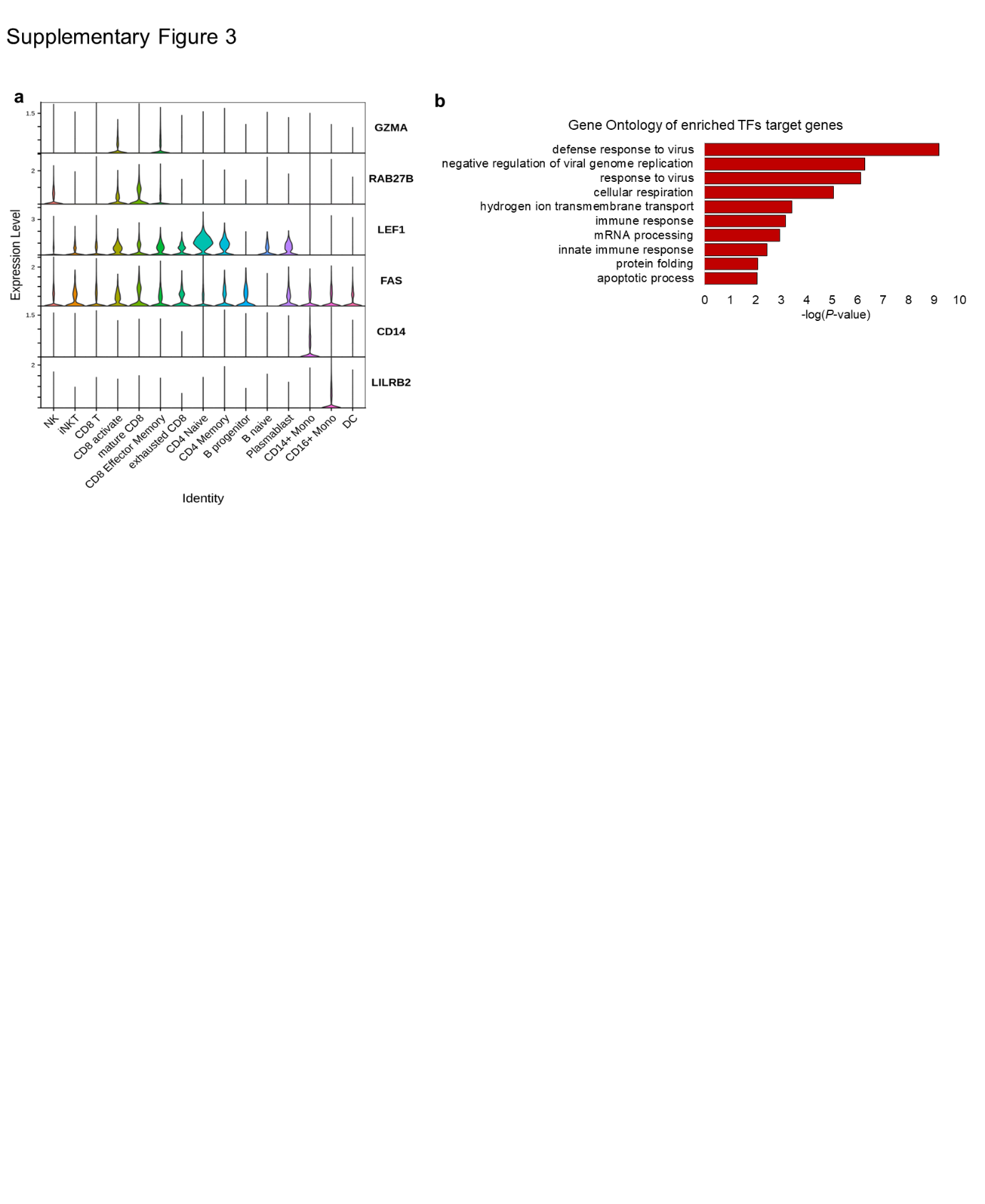
**Fig. S3**. **Single-cell multi-omics analysis identifies epigenetic characteristics of scATAC-seq dataset and upregulated transcription factors in HIV-1 RNA+ cells**

**(A)** Violin plot displays the gene expression of marker genes in scATAC-seq dataset.

**(B)** The bar plot represents the gene ontologies associated with the target genes of the upregulated TFs in HIV-1 RNA+ cells (*P*-value < 0.01). The significance of these enrichments is expressed as -log(*P*-value).

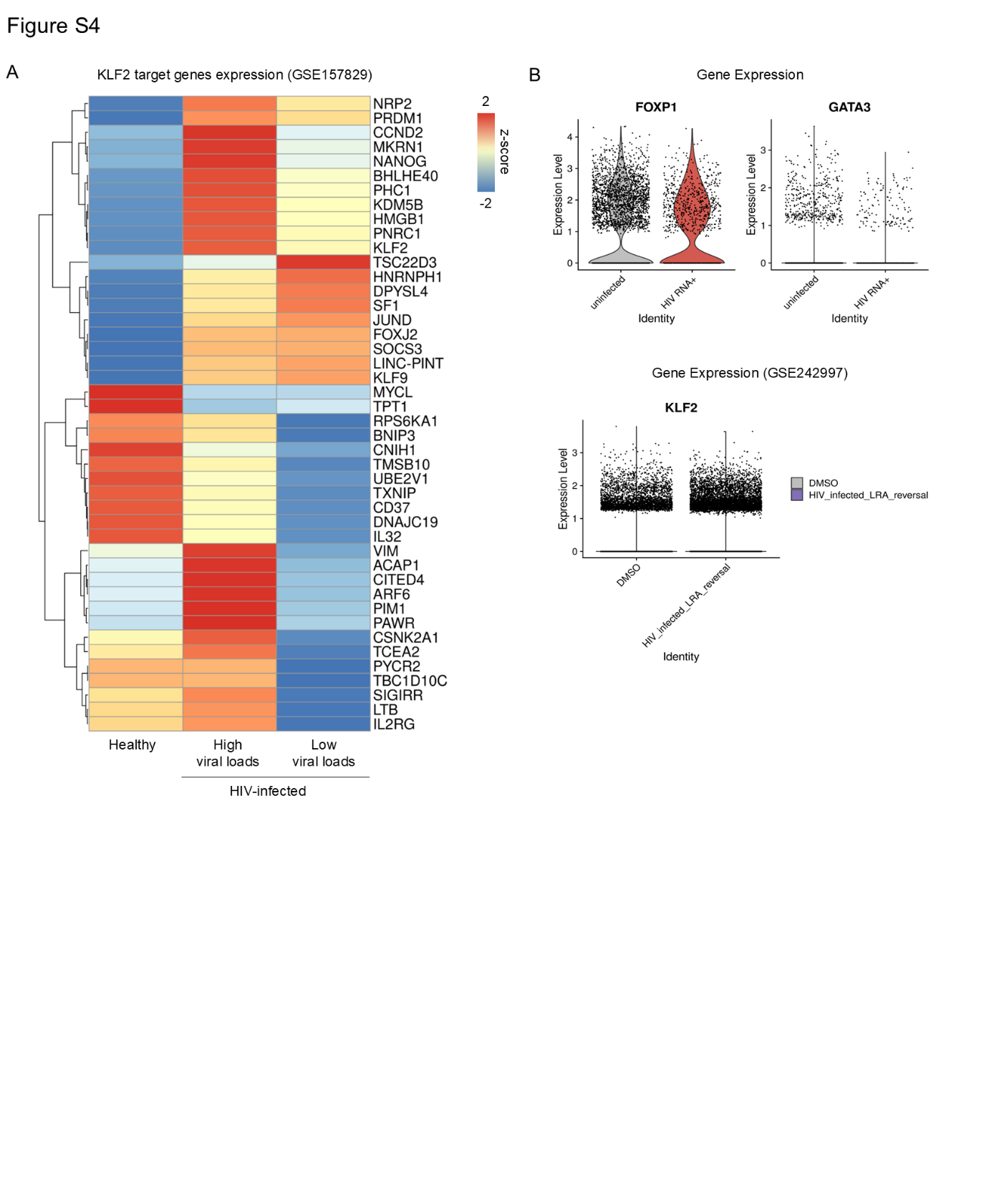

**Fig. S4**. **KLF2 target gene expression in HIV-1-infected CD4+ T cells**

**(A)** Heatmap showing the expression (Z-score normalized) of KLF2 target genes across healthy controls and HIV-1-infected individuals with high and low viral loads. Gene expression data were obtained from a published dataset (GSE157829)

**(B)** Violin plots showing the expression levels of *FOXP1* and *GATA3* in HIV-1 RNA+ versus uninfected CD4+ T cells (top, current study). Bottom panel shows expression levels of *KLF2* in published dataset (GSE242997), comparing HIV-infected cells treated with latency-reversing agents (LRA) and DMSO controls.

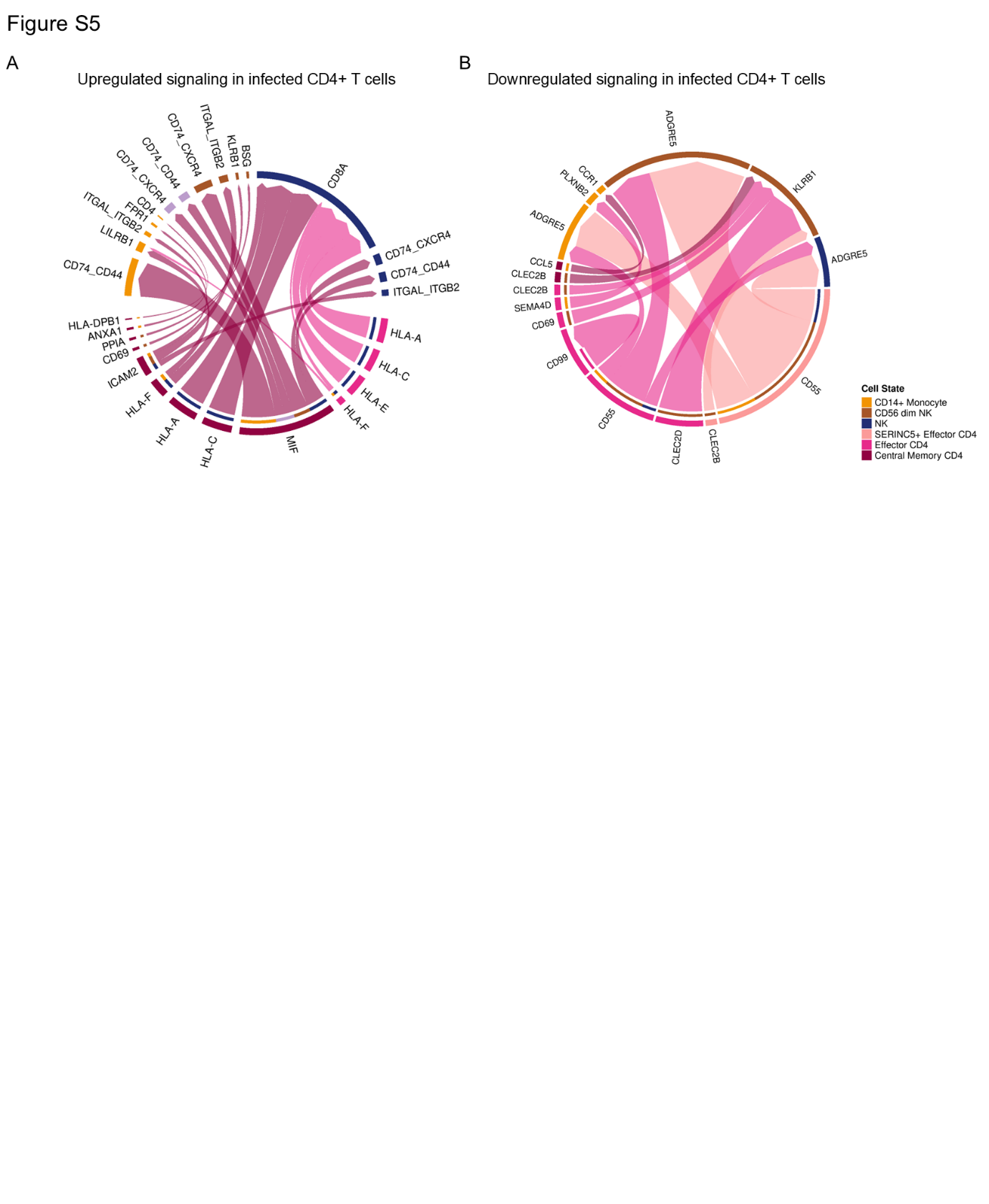

**Fig. S5**. **Cell-cell interaction in HIV-1-infected CD4+ T cells**

(A) Circos plot show cell–cell interactions that are upregulated in HIV-1-infected CD4+ T cells. Line thickness represents interaction strength, and arcs are colored by cell types.
(B) Circos plot shows downregulated interactions in HIV-1 infected CD4+ T cells.

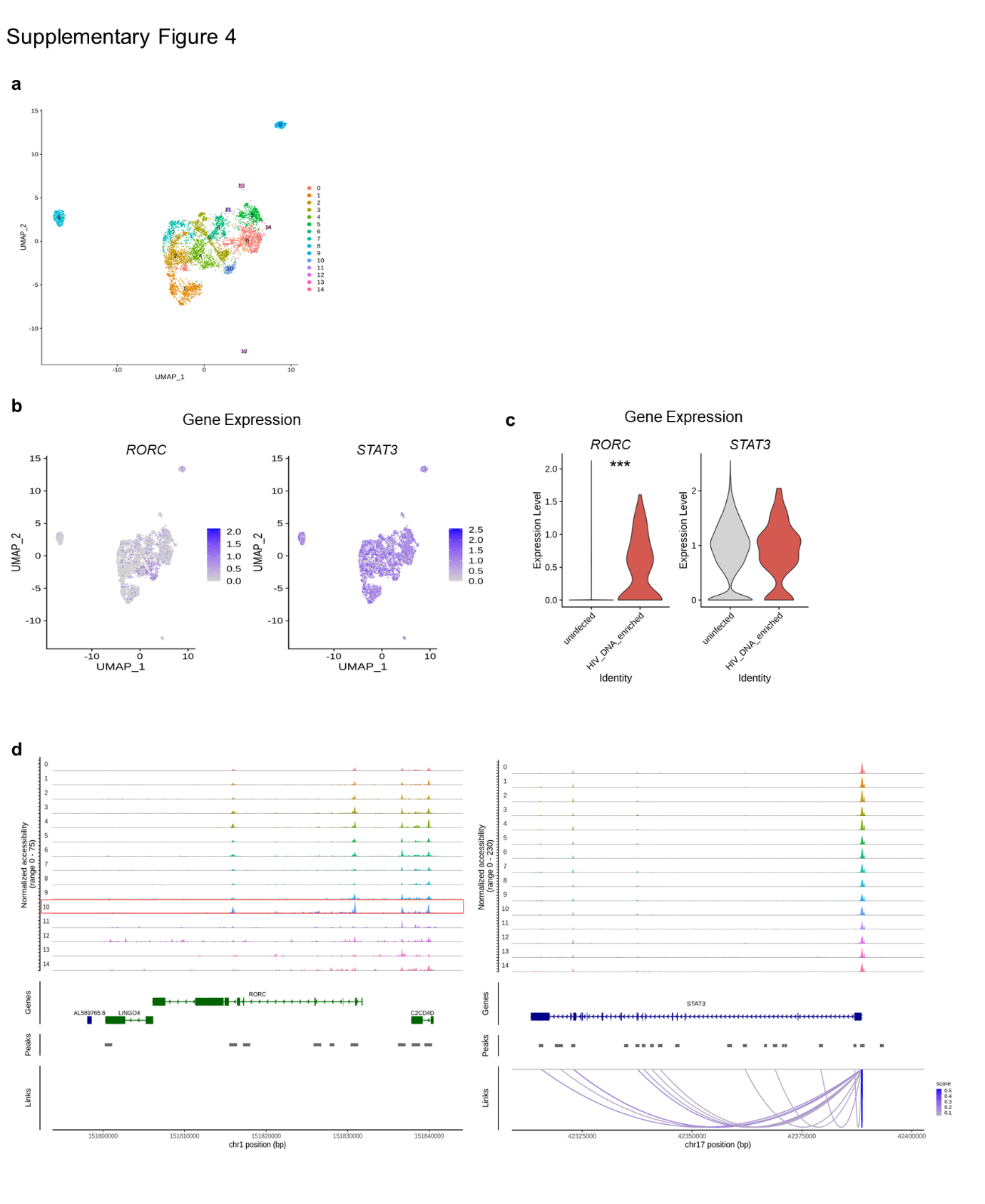
**Fig. S6**. **Single-cell epigenetic analysis identifies HIV-1 DNA+ cells are mostly represented by Th17 cell type.**

**(A)** The ATAC UMAP plot displays the distribution of 5,608 CD4 T cells from early infected patients.

**(B-C)** The Feature Plot and violin plot indicate gene expression (*RORC*, *STAT3*) in CD4 T cells. **P* < 0.01

**(D)** The tracks display the normalized chromatin accessibility levels at the promoter regions of *RORC* and *STAT3* in all cluster of CD4 T cells.

**Supplementary Tables**

| Sample ID | Sex | Age | Duration from HIV infection (Day) | CD4+ T  (cells/mm3) | HIV-1  Viral loads  (Log10 copies/ml) | HIV-1 p24 Antigen | HIV-1  Antibodies  (PA) | Sequencing  Techniques | | |
| --- | --- | --- | --- | --- | --- | --- | --- | --- | --- | --- |
|  |  |  |  |  |  |  |  | scRNA-seq | scATAC-seq | sc-Multiome |
| 06-D10629 | M | 51 | 158 | 760 | 16.1 | Positive | Negative | No | No | Yes |
| C1-C80404 | M | 17 | 0 | 443 | 13.4 | Positive | Negative | Yes | No | Yes |
| 35-C90552 | M | 45 | 20 | 655 | 13.2 | Positive | Negative | Yes | No | Yes |
| 36-C70518 | M | 28 | 3 | N/A | 14.9 | N/A | N/A | Yes | No | Yes |
| 02-C80622 | M | 28 | 0 | 970 | 16.1 | Positive | Negative | Yes | Yes | No |
| 02-D00365 | M | 34 | 12 | 482 | 15 | Negative | Negative | Yes | Yes | No |
| 36-C80800 | F | 50 | 0 | 643 | 16.1 | Positive | Negative | Yes | Yes | No |
| 36-D00564 | M | 50 | 175 | 888 | 13.3 | Negative | Positive | Yes | Yes | No |
| 02-C80649 | M | 59 | 0 | 597 | 15.7 | N/A | N/A | Yes | No | No |

* M: Male, F: Female

**Table S1. Epidemiological and Virological Characteristics, and Sequencing Data Information for 9 Acute HIV-infected Patients at Baseline**

| **Gene** | **p_val** | **avg_logFC** | **pct.1** | **pct.2** | **p_val_adj** |
| --- | --- | --- | --- | --- | --- |
| TPT1 | 2.17E-74 | 0.664730297 | 0.887 | 0.829 | 6.97E-70 |
| VIM | 4.75E-44 | 0.486947637 | 0.711 | 0.507 | 1.52E-39 |
| TMSB10 | 1.01E-67 | 0.484596487 | 0.924 | 0.821 | 3.25E-63 |
| CD37 | 7.61E-57 | 0.483621046 | 0.615 | 0.344 | 2.44E-52 |
| HMGB1 | 1.26E-55 | 0.449454734 | 0.786 | 0.554 | 4.05E-51 |
| LTB | 4.33E-46 | 0.44052761 | 0.839 | 0.644 | 1.39E-41 |
| TXNIP | 2.55E-64 | 0.408496641 | 0.955 | 0.843 | 8.16E-60 |
| IL32 | 2.84E-29 | 0.400628464 | 0.723 | 0.559 | 9.10E-25 |
| KLF2 | 2.62E-45 | 0.383897324 | 0.777 | 0.546 | 8.41E-41 |
| PNRC1 | 1.45E-30 | 0.299190983 | 0.651 | 0.434 | 4.64E-26 |
| ACAP1 | 5.87E-25 | 0.270001786 | 0.553 | 0.358 | 1.88E-20 |
| IL2RG | 2.18E-25 | 0.259545545 | 0.413 | 0.233 | 7.00E-21 |
| SF1 | 3.40E-16 | 0.211089385 | 0.8 | 0.639 | 1.09E-11 |
| TBC1D10C | 1.01E-15 | 0.208888529 | 0.382 | 0.244 | 3.23E-11 |
| PIM1 | 2.00E-14 | 0.207804659 | 0.361 | 0.233 | 6.42E-10 |
| SIGIRR | 3.57E-13 | 0.198079587 | 0.321 | 0.203 | 1.14E-08 |
| CCND2 | 5.82E-12 | 0.189860451 | 0.311 | 0.198 | 1.87E-07 |
| HNRNPH1 | 1.00E-15 | 0.188659263 | 0.776 | 0.62 | 3.22E-11 |
| TSC22D3 | 9.49E-15 | 0.171356073 | 0.467 | 0.317 | 3.04E-10 |
| DNAJC19 | 3.04E-17 | 0.167563816 | 0.198 | 0.095 | 9.74E-13 |
| CITED4 | 3.31E-16 | 0.1675141 | 0.206 | 0.103 | 1.06E-11 |
| UBE2V1 | 1.21E-14 | 0.14695742 | 0.121 | 0.05 | 3.89E-10 |
| JUND | 1.38E-15 | 0.140976983 | 0.559 | 0.383 | 4.43E-11 |
| MKRN1 | 1.04E-05 | 0.13071128 | 0.245 | 0.179 | 0.333091 |
| PRDM1 | 2.48E-05 | 0.122195235 | 0.117 | 0.074 | 0.796627 |
| BNIP3 | 3.35E-11 | 0.116108557 | 0.119 | 0.056 | 1.07E-06 |
| LINC-PINT | 0.008899725 | 0.071251147 | 0.394 | 0.341 | 1 |
| TCEA2 | 0.000497195 | 0.068589737 | 0.048 | 0.026 | 1 |
| PYCR2 | 2.40E-07 | 0.060669396 | 0.119 | 0.067 | 0.00771 |
| KLF9 | 0.001952912 | 0.058854945 | 0.21 | 0.163 | 1 |
| CNIH1 | 1.34E-07 | 0.056829405 | 0.125 | 0.07 | 0.004296 |
| ARF6 | 0.000182984 | 0.054848756 | 0.208 | 0.154 | 1 |
| BHLHE40 | 0.000411762 | 0.053459996 | 0.058 | 0.033 | 1 |
| RPS6KA1 | 0.024616225 | 0.044512378 | 0.088 | 0.066 | 1 |
| CSNK2A1 | 0.002730067 | 0.038988524 | 0.189 | 0.146 | 1 |
| PAWR | 0.000829052 | 0.036027924 | 0.018 | 0.007 | 1 |
| FOXJ2 | 0.830108419 | 0.025027762 | 0.023 | 0.025 | 1 |
| SOCS3 | 0.012105249 | 0.017691013 | 0.143 | 0.111 | 1 |
| NRP2 | 0.188390647 | 0.010845121 | 0.003 | 0.001 | 1 |
| DPYSL4 | 0.001187095 | 0.010504532 | 0.003 | 0 | 1 |
| KDM5B | 0.355956849 | 0.006549433 | 0.132 | 0.12 | 1 |
| NANOG | 0.181105411 | 0.005793679 | 0.002 | 0.001 | 1 |
| PHC1 | 0.69964956 | 0.005013823 | 0.035 | 0.032 | 1 |
| MYCL | 0.344726739 | 0.000883179 | 0.001 | 0 | 1 |
| S1PR3 | 0.744460326 | -0.000627781 | 0.001 | 0.001 | 1 |
| DNMT3A | 0.489967984 | -0.001010646 | 0.141 | 0.15 | 1 |
| RRP1B | 0.205136712 | -0.001104724 | 0.163 | 0.143 | 1 |
| NXNL2 | 0.450035052 | -0.001541665 | 0 | 0.001 | 1 |
| DMRT1 | 0.450035052 | -0.001673579 | 0 | 0.001 | 1 |
| BMP7 | 0.450035052 | -0.001896741 | 0 | 0.001 | 1 |
| SNAI1 | 0.450035052 | -0.001985345 | 0 | 0.001 | 1 |
| MFSD2A | 0.354768244 | -0.002548496 | 0 | 0.001 | 1 |
| ZMYM3 | 0.958216028 | -0.003288628 | 0.02 | 0.021 | 1 |
| PLA2G1B | 0.450035052 | -0.003297112 | 0 | 0.001 | 1 |
| NOTCH3 | 0.450035052 | -0.004169338 | 0 | 0.001 | 1 |
| TRPM3 | 0.613033626 | -0.006833198 | 0.001 | 0.002 | 1 |
| RAD51C | 0.052466005 | -0.010661489 | 0.053 | 0.038 | 1 |
| CDKN1A | 0.090790919 | -0.013386654 | 0 | 0.003 | 1 |
| DENND2A | 0.054112543 | -0.014882968 | 0.001 | 0.006 | 1 |
| HOXB4 | 0.634292218 | -0.017283224 | 0.015 | 0.017 | 1 |
| RABIF | 0.486560265 | -0.025469725 | 0.029 | 0.034 | 1 |
| RPN1 | 0.438232729 | -0.026026513 | 0.119 | 0.108 | 1 |
| KLF4 | 0.157228999 | -0.028322843 | 0 | 0.002 | 1 |
| AP1M1 | 0.66257167 | -0.032616089 | 0.093 | 0.098 | 1 |
| HOOK2 | 0.214501657 | -0.033820571 | 0.056 | 0.066 | 1 |
| PGS1 | 0.765659484 | -0.056894937 | 0.061 | 0.057 | 1 |
| TGIF1 | 0.68822548 | -0.06021225 | 0.035 | 0.037 | 1 |
| AP5M1 | 0.282209538 | -0.061633796 | 0.09 | 0.101 | 1 |
| SP1 | 0.372255551 | -0.069005677 | 0.057 | 0.064 | 1 |
| ATG4C | 0.014615343 | -0.080024748 | 0.021 | 0.037 | 1 |
| PXN | 0.083160525 | -0.09286964 | 0.118 | 0.137 | 1 |
| DLGAP1 | 0.000585314 | -0.110979077 | 0.003 | 0.018 | 1 |
| TEX14 | 3.59E-08 | -0.127037624 | 0.005 | 0.041 | 0.001152 |
| CD83 | 0.000206579 | -0.129834592 | 0.006 | 0.025 | 1 |
| CYLD | 0.003516763 | -0.141271159 | 0.531 | 0.546 | 1 |
| SMARCAD1 | 0.000407205 | -0.201335112 | 0.066 | 0.102 | 1 |
| MSRA | 0.000171836 | -0.232231502 | 0.059 | 0.095 | 1 |
| FOS | 0.000404187 | -0.233427767 | 0.032 | 0.061 | 1 |
| RAD23B | 6.14E-05 | -0.238391059 | 0.135 | 0.182 | 1 |
| ARL8B | 2.25E-06 | -0.24180768 | 0.08 | 0.133 | 0.072268 |
| JUNB | 0.000379409 | -0.274886806 | 0.504 | 0.519 | 1 |
| JARID2 | 6.74E-08 | -0.291366599 | 0.148 | 0.221 | 0.002162 |
| RYBP | 3.05E-10 | -0.329940877 | 0.062 | 0.133 | 9.78E-06 |
| ATG5 | 4.10E-10 | -0.377720823 | 0.105 | 0.18 | 1.32E-05 |
| SLC2A3 | 2.14E-11 | -0.406462164 | 0.143 | 0.235 | 6.87E-07 |
| PPP1R12A | 2.37E-14 | -0.415261823 | 0.319 | 0.426 | 7.60E-10 |
| DTNB | 4.43E-21 | -0.496467883 | 0.044 | 0.158 | 1.42E-16 |
| KANK1 | 5.65E-20 | -0.543001768 | 0.042 | 0.15 | 1.81E-15 |

**Table S2. Differentially expressed genes overlapped with KLF2 target gene in CD4 T cell. (HIV-1 RNA+ vs. Uninfected)**

| Primer name | sequence (5'-3') |
| --- | --- |
| P1 | GAAATCTGTTGACTCAGATTGGTTGCACTTTAAATTTTCCCATTAGCC |
| P2 | CTATGGCAGGAAGAAGCGGAGACAGCGACGAAGAGCTCCTCA |
| P3 | CTACAAGGGACTTTCCGCTGGGGACTTTCCAGGGAGGCGTGG |

**Table S3. Primers used for HIV-1 targeted sequencing**
